# Mutations that improve the efficiency of a weak-link enzyme are rare compared to adaptive mutations elsewhere in the genome

**DOI:** 10.1101/624205

**Authors:** Andrew B. Morgenthaler, Wallis R. Kinney, Christopher C. Ebmeier, Corinne M. Walsh, Daniel J. Snyder, Vaughn S. Cooper, William M. Old, Shelley D. Copley

## Abstract

New enzymes often evolve by amplification and divergence of genes encoding enzymes with a weak ability to provide a new function. Experimental studies to date have followed the evolutionary trajectory of an amplified gene, but have not addressed other mutations in the genome when fitness is limited by an evolving gene. We have adapted *Escherichia coli* in which an enzyme’s weak secondary activity has been recruited to serve an essential function. While the gene encoding the “weak-link” enzyme amplified in all eight populations, mutations improving the new activity occurred in only one. This beneficial allele quickly swept the amplified array, displacing the parental allele. Most adaptive mutations, however, occurred elsewhere in the genome. We have identified the mechanisms by which three of the classes of mutations increase fitness. These mutations may be detrimental once a new enzyme has evolved, and require reversion or compensation, leading to permanent changes in the genome.

## Introduction

The expansion of huge superfamilies of enzymes, transcriptional regulators, transporters, and signaling molecules from single ancestral genes has been a dominant process in the evolution of life (Bergthorsson et al., 2007; Chothia et al., 2003; Glasner et al., 2006; A. L. Hughes, 1994; Ohno, 1970; Todd et al., 2001). The emergence of new protein family members has enabled organisms to access new nutrients, sense new stimuli, and respond to changing conditions with ever more sophistication (Bouquin & Khila, 2017; Conant & Wolfe, 2008; Nei & Rooney, 2005; Reams & Neidle, 2004; Starr et al., 2017; Storz, 2016).

The <u>**I**</u>nnovation-<u>**A**</u>mplification-<u>**D**</u>ivergence (IAD) model (Figure 1) posits that evolution of new genes by duplication and divergence begins when a physiologically irrelevant side activity of a protein, known as a promiscuous activity (Copley, 2003, 2017), becomes important for fitness due to a mutation or environmental change (Bergthorsson et al., 2007; Francino, 2005; A. L. Hughes, 1994; Näsvall et al., 2012). Often this newly useful activity is inefficient, making the enzyme the “weak-link” in metabolism. Gene duplication/amplification provides a ready mechanism to improve fitness by increasing the abundance of a weak-link enzyme. If further mutations lead to evolution of an enzyme capable of efficiently carrying out the newly needed function, selective pressure to maintain a high copy number will be removed, allowing extra copies to be lost and leaving behind two paralogs (or just one gene encoding a new enzyme if the original function is no longer needed).

**Figure 1.**
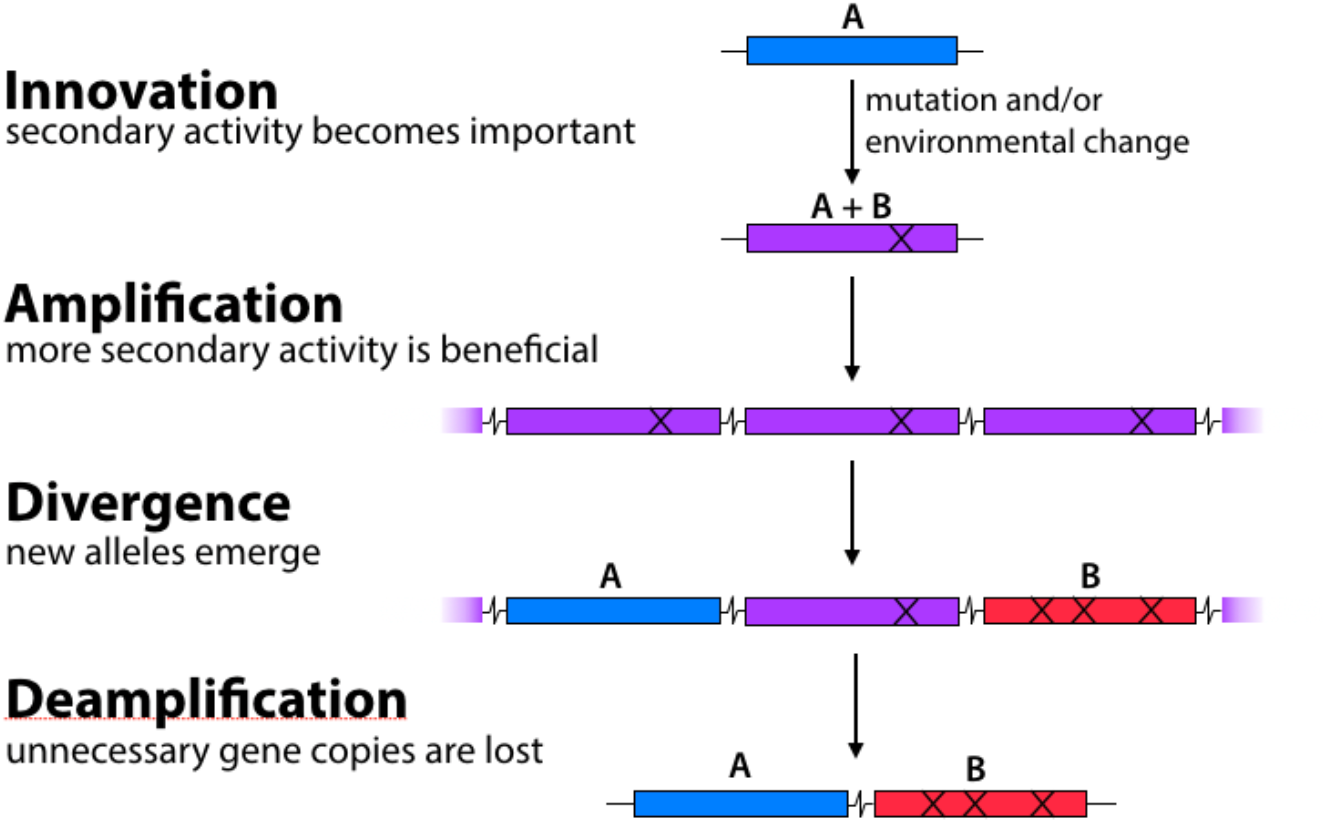
The Innovation-Amplification-Divergence (IAD) model of gene evolution. A physiologically irrelevant side activity B of an enzyme may become physiologically relevant due to a mutation or environmental change. Gene amplification increases the abundance of the weak-link enzyme. Acquisition of mutations can improve the efficiency of the newly important activity B. Once sufficient B activity is achieved, selection is relaxed and extra gene copies are lost, leaving behind two paralogs.

While the IAD model provides a satisfying theoretical framework for the process of gene duplication and divergence, our understanding of the process is far from perfect. Although the signatures of gene duplication and divergence are obvious in extant genomes, we have little information about the genome contexts and environments in which new enzymes arose. Laboratory evolution offers the possibility of tracking this process in real time. In a landmark study, Näsvall et al. used laboratory evolution to demonstrate that a gene encoding an enzyme with two inefficient activities required for synthesis of histidine and tryptophan amplified and diverged to alleles encoding two specialists within 2000 generations (Näsvall et al., 2012). However, this study followed only mutations in the diverging gene. In general, when an organism is exposed to a novel selection pressure that requires evolution of a new enzyme, any mutation – either in the gene encoding the weak-link enzyme or elsewhere in the genome – that improves fitness will provide a selective advantage.

We have generated a model system in *E. coli* for exploring the relative importance of mutations in a gene encoding a weak-link enzyme and elsewhere in the genome. ProA (γ-glutamyl phosphate reductase, Figure 2) is essential for proline synthesis in *E. coli*. ArgC (*N*-acetylglutamyl phosphate reductase) catalyzes a similar reaction in the arginine synthesis pathway, although the two enzymes are not homologous (Goto et al., 2003; Ludovice et al., 1992; Page et al., 2003). ProA can reduce *N*-acetylglutamyl phosphate, but its activity is too inefficient to support growth of a *∆argC* strain of *E. coli* in glucose. However, a point mutation that changes Glu383 to Ala allows slow growth of the *∆argC* strain in glucose. Enzymatic assays show that E383A ProA (ProA*) has severely reduced activity with γ-glutamyl semialdehyde (GSA), but substantially improved activity with *N*-acetylglutamyl semialdehyde (NAGSA) (Khanal et al., 2015; McLoughlin & Copley, 2008). (It is necessary to assay kinetic parameters in the reverse direction because the substrates for the forward reaction are too unstable to prepare and purify.) Glu383 is in the active site of the enzyme; the change to Ala may create extra room in the active site to accommodate the larger substrate for ArgC, but at a cost to the ability to bind and orient the native substrate. The poor efficiency of the weak-link ProA* creates strong selective pressure for improvement of both proline and arginine synthesis during growth of *∆argC E. coli* on glucose as a sole carbon source.

**Figure 2.**
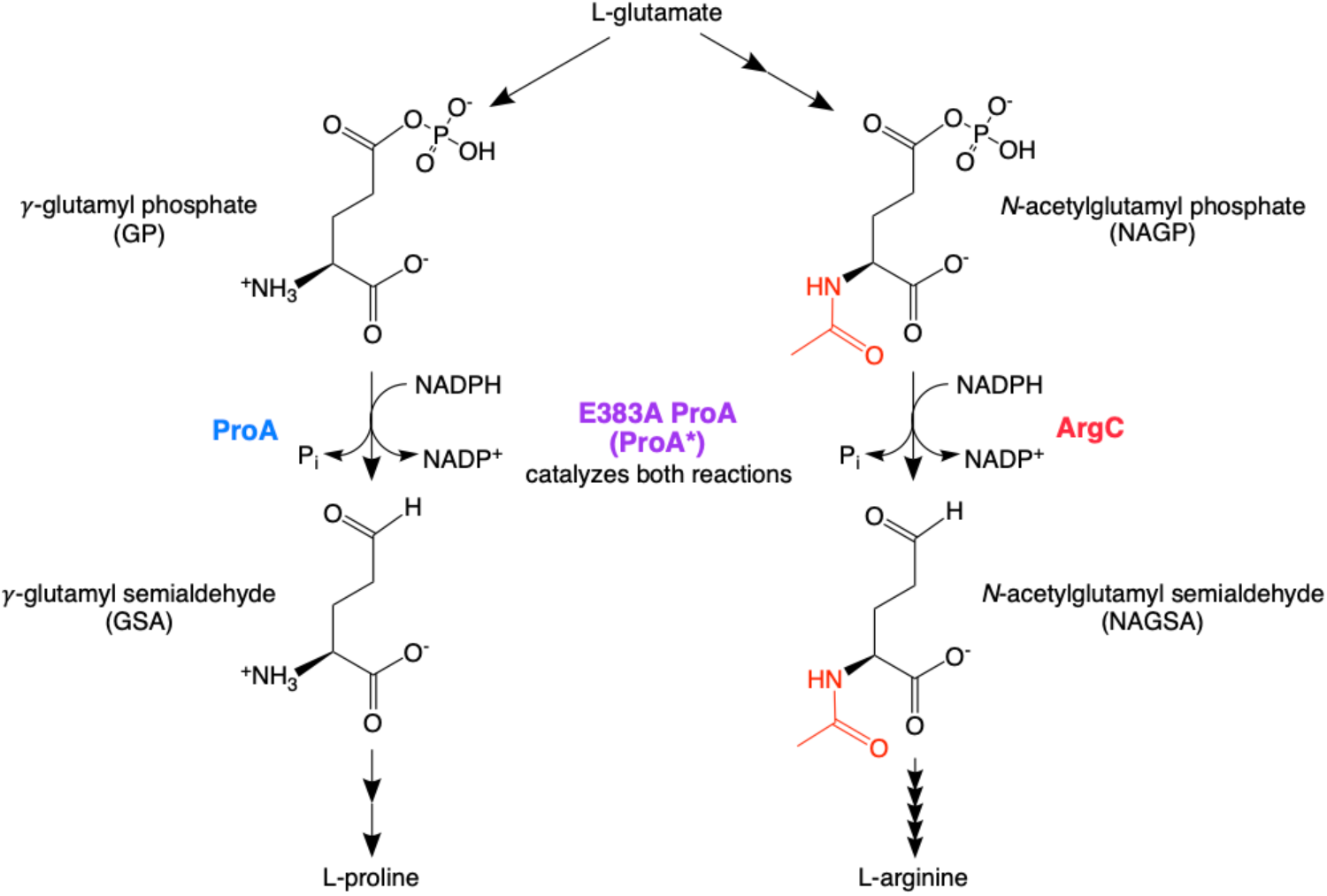
Reactions catalyzed by ProA (γ-glutamyl phosphate reductase) in proline synthesis and ArgC (*N*-acetylglutamyl phosphate reductase) in arginine synthesis. E383A ProA (ProA*) can catalyze both reactions, albeit poorly.

We adapted eight replicate populations of *∆argC proA* E. coli* in minimal medium supplemented with glucose and proline for up to 1000 generations to identify mechanisms by which the impairment in arginine synthesis could be alleviated. Our expectation that amplification of *proA\** would be beneficial was borne out in all populations. However, whole-genome sequencing of the adapted populations showed that a mutation in *proA\** occurred in only one population. Indeed, most of the adaptive mutations occurred outside of *proA\**. We have identified the mechanisms by which three common classes of mutations allow a relatively rapid increase in fitness while the more difficult process of improving the weak-link enzyme progresses. However, they increase fitness at a cost to presumably well-evolved functions.

Our results demonstrate that mutations elsewhere in the genome play an important role during the process of gene amplification and divergence when the inefficient activity of a weak-link enzyme limits fitness. Thus, the process of evolution of a new enzyme by gene duplication and divergence is inextricably intertwined with mutations elsewhere in the genome that improve fitness by other mechanisms.

## Results

### Growth rate of *∆argC proA* E. coli* increased 3-fold within a few hundred generations of adaptation in M9/glucose/proline

We generated a progenitor strain for laboratory evolution by replacing *argC* with the *kan*^*r*^ antibiotic resistance gene, modifying *proA* to encode ProA*, and introducing a mutation in the - 10 region of the promoter of the *proBA* operon that was previously shown to occur frequently during adaptation of the *∆argC* strain (Kershner et al., 2016). We also introduced *yfp* downstream of *proA\** and deleted several genes (*fimAICDFGH* and *csgBAC*, which are required for the formation of fimbriae and curli, respectively (Barnhart & Chapman, 2006; Proft & Baker, 2009)) to minimize the occurrence of biofilms. We evolved eight parallel lineages of this strain (AM187, Table 1) in M9 minimal medium supplemented with 0.2% glucose, 0.4 mM proline, and 20 µg/mL kanamycin in a turbidostat to identify mutations that improve synthesis of arginine. We used a turbidostat rather than a serial transfer protocol because turbidostats can maintain cultures in exponential phase and thereby avoid selection for mutations that simply decrease lag phase or improve survival in stationary phase. Turbidostats also avoid population bottlenecks during serial passaging that can result in loss of genetic diversity.

**Table 1.**
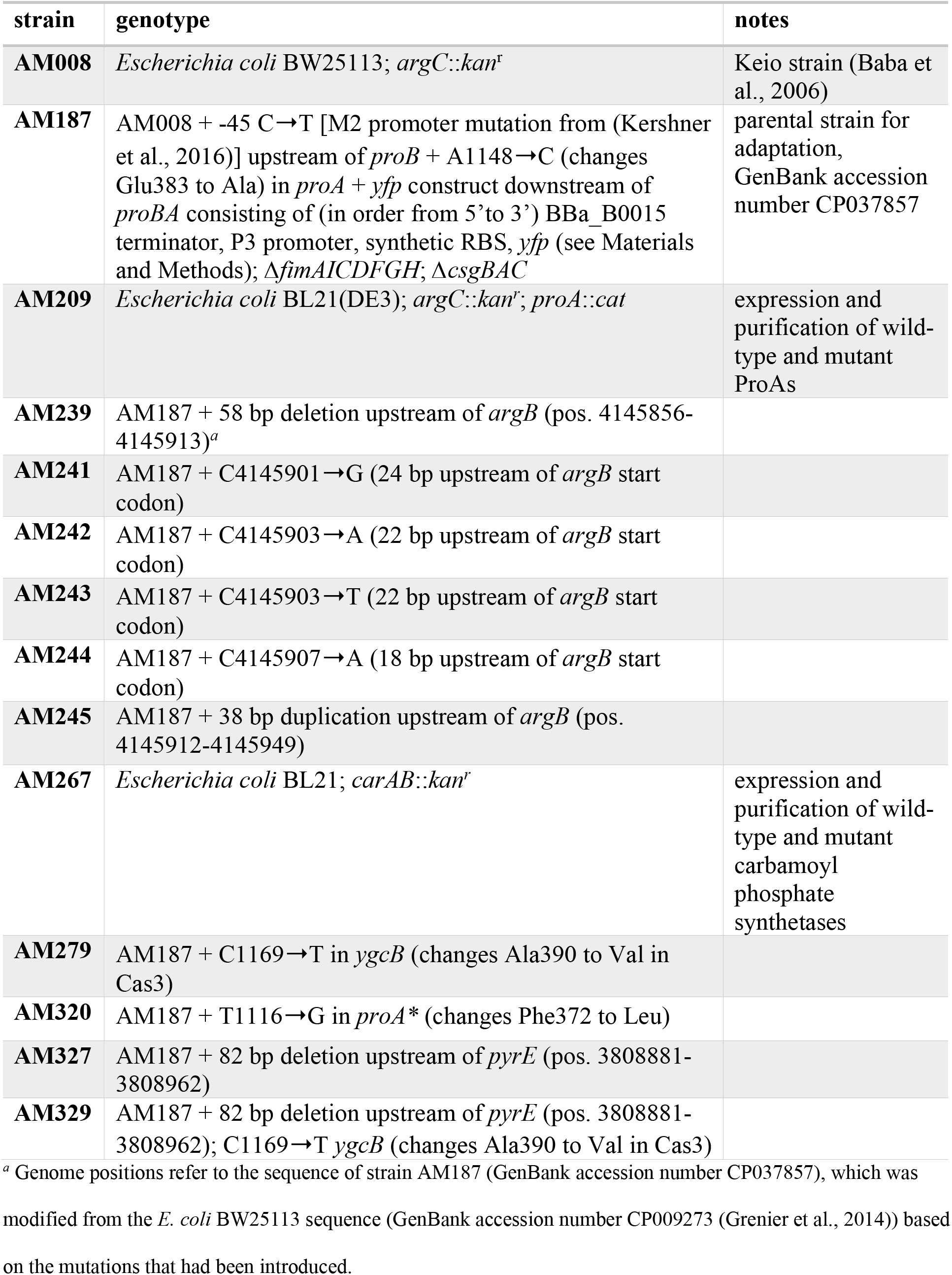
Strains used in this work.

| strain | genotype | notes |
| --- | --- | --- |
| **AM008** | *Escherichia coli* BW25113; *argC::kan<sup>r</sup>* | Keio strain (Baba et al., 2006) |
| **AM187** | AM008 + -45 C→T [M2 promoter mutation from (Kershner et al., 2016)] upstream of *proB* + A1148→C (changes Glu383 to Ala) in *proA* + *yfp* construct downstream of *proBA* consisting of (in order from 5' to 3') BBa_B0015 terminator, P3 promoter, synthetic RBS, *yfp* (see Materials and Methods); $\Delta fimAICDFGH$; $\Delta csgBAC$ | parental strain for adaptation, GenBank accession number CP037857 |
| **AM209** | *Escherichia coli* BL21(DE3); *argC::kan<sup>r</sup>*; *proA::cat* | expression and purification of wild-type and mutant ProAs |
| **AM239** | AM187 + 58 bp deletion upstream of *argB* (pos. 4145856-4145913)<sup>a</sup> |  |
| **AM241** | AM187 + C4145901→G (24 bp upstream of *argB* start codon) |  |
| **AM242** | AM187 + C4145903→A (22 bp upstream of *argB* start codon) |  |
| **AM243** | AM187 + C4145903→T (22 bp upstream of *argB* start codon) |  |
| **AM244** | AM187 + C4145907→A (18 bp upstream of *argB* start codon) |  |
| **AM245** | AM187 + 38 bp duplication upstream of *argB* (pos. 4145912-4145949) |  |
| **AM267** | *Escherichia coli* BL21; *carAB::kan<sup>r</sup>* | expression and purification of wild-type and mutant carbamoyl phosphate synthetases |
| **AM279** | AM187 + C1169→T in *ygcB* (changes Ala390 to Val in Cas3) |  |
| **AM320** | AM187 + T1116→G in *proA* * (changes Phe372 to Leu) |  |
| **AM327** | AM187 + 82 bp deletion upstream of *pyrE* (pos. 3808881-3808962) |  |
| **AM329** | AM187 + 82 bp deletion upstream of *pyrE* (pos. 3808881-3808962); C1169→T *ygcB* (changes Ala390 to Val in Cas3) |  |

<sup>a</sup> Genome positions refer to the sequence of strain AM187 (GenBank accession number CP037857), which was modified from the *E. coli* BW25113 sequence (GenBank accession number CP009273 (Grenier et al., 2014)) based on the mutations that had been introduced.

Growth rate in each culture tube was averaged over each 24-hour period and was used to calculate the number of generations each day. Each culture was maintained until a biofilm formed (33-57 days, corresponding to 470-1000 generations). While it is possible to restart cultures from individual clones after biofilm formation, this practice introduces a severe population bottleneck. Thus, we decided to stop the adaptation for each population when a biofilm formed. For this reason, every population was adapted for a different number of generations.

Over the course of the experiment, growth rate increased 2.5-3.5-fold for all eight populations (Figure 3). This improvement corresponds to a change in doubling time from ~3.3 hours to ~1 hour. As expected, the rate at which growth rate increased slowed over the course of adaptation. Occasional dips in growth rate occurred during the adaptation (Figure 3). These dips are artifacts arising from temporary aberrations in selective conditions due to turbidostat malfunctions that prevented introduction of fresh medium, causing the cultures to enter stationary phase. Occasionally cultures were saved as frozen stocks until the turbidostat was fixed (see Materials and Methods). Restarting cultures from frozen stocks may have caused a temporary drop in growth rate.

**Figure 3.**
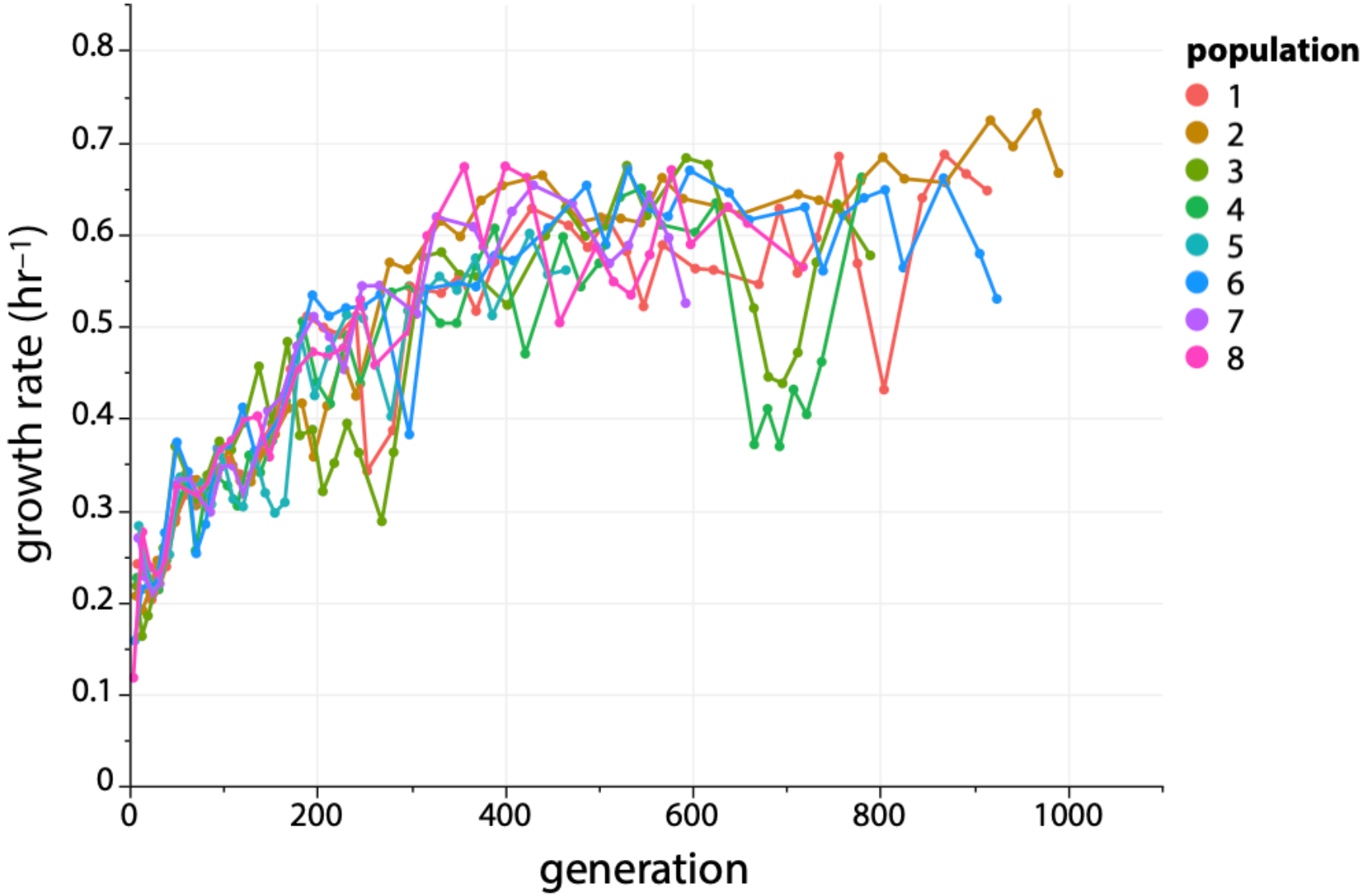
Growth rate increases ~3-fold during adaptation of *∆argC* M2-*proA* E. coli* in M9 minimal medium containing 0.2% glucose, 0.4 mM proline and 20 µg/mL kanamycin.

### The copy number and extent of amplification around *proA\** varied among replicate populations

We monitored *proA\** copy number during the adaptation experiment using qPCR of population genomic DNA (Figure 4A). *proA*\* was present in at least 6 copies by generation 300 in all eight populations. Six of the populations maintained 6-9 copies for the remainder of the adaptation. In contrast, *proA\** copy number in population 2 increased to as many as 20 copies. In population 3, *proA\** copy number dropped to three by generation 400.

**Figure 4.**
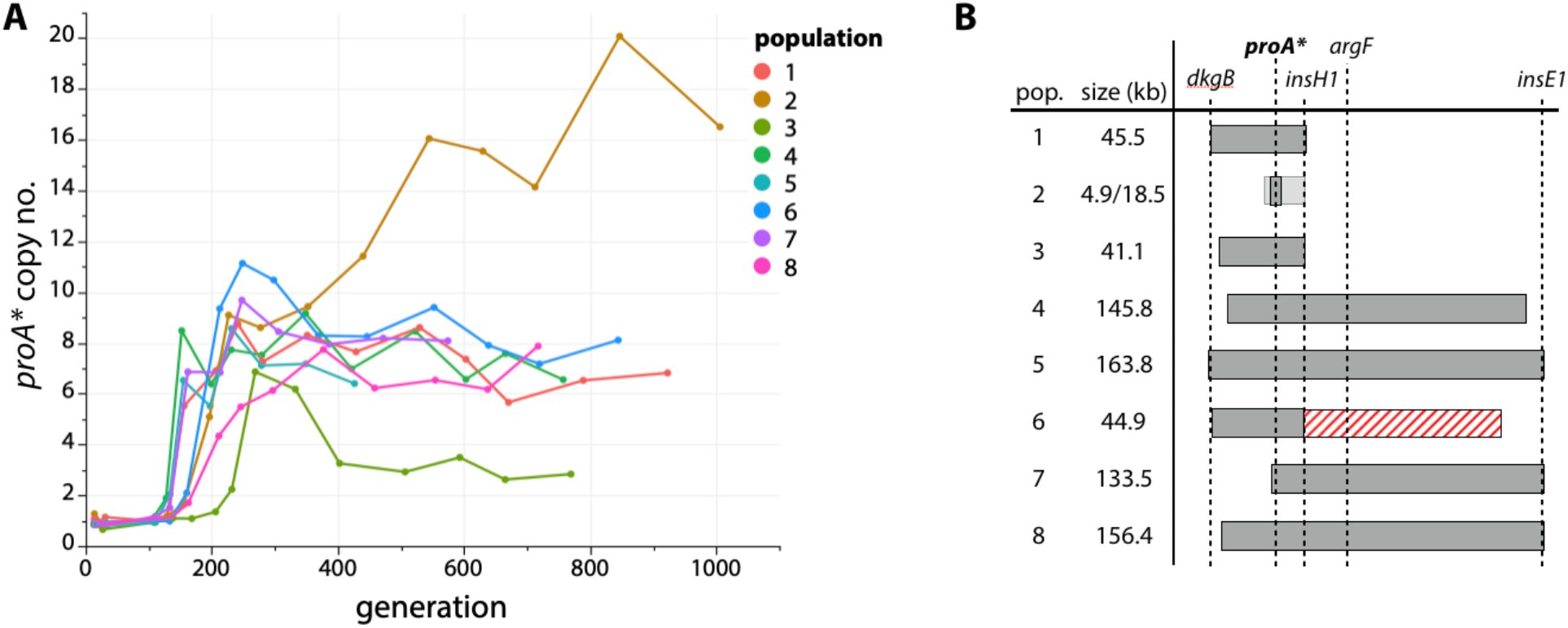
*proA\** is amplified during adaptation. (A) *proA\** copy number in each adapted population as measured by qPCR. (B) Regions of genomic amplification in each adapted population based population-wide, whole-genome sequencing. Population 2 had two overlapping regions of amplification, both of which included *proA\** (shown as differently shaded bars). Population 6 had a 95.1 kb deletion (shown as a red striped bar) immediately downstream of the amplified region.

We identified the boundaries of the amplified regions in all eight populations by sequencing population genomic DNA (Figure 4B, Supplementary File 1). The amplified region in population 2 was unusually small, spanning only 4.9 kb and resulting in co-amplification of only two other genes besides *proBA\**. Population 2 also appeared to have a second region of amplification of 18.5 kb. (Whether these two distinct amplification regions coexisted in the same clone or as two separate clades within the population could not be determined from population genome sequencing.) In contrast, the amplified regions in the other seven populations ranged from 41.1 to 163.8 kb, encompassing between 55 and 177 genes. We attribute the variation in *proA\** copy number to these differences in the size of the amplified region on the genome. The population with the smallest amplified region (4.9 kb, population 2) carries fewer multicopy genes and thus should incur a lower fitness cost, allowing *proA\** to reach a higher copy number (Adler et al., 2014; Kugelberg et al., 2006; Pettersson et al., 2009; Reams et al., 2010).

### A mutation in *proA\** led to deamplification in population 3

The decrease in *proA\** copy number in population 3 was particularly noteworthy since it might have been an indication that a mutation had improved the neo-ArgC activity of ProA*, resulting in a decreased need for multiple copies. In fact, a mutation in *proA\** that changes Phe372 to Leu (Figure 5A) was observed in population 3. E383A F372L ProA will be designated ProA** hereafter. Introduction of this mutation into the parental strain (which carried *proA\**) increased growth rate by 69% (Figure 5B), confirming that the mutation is adaptive and not a random hitchhiker. In contrast, no mutations in *proA\** were identified in any of the other populations.

**Figure 5.**
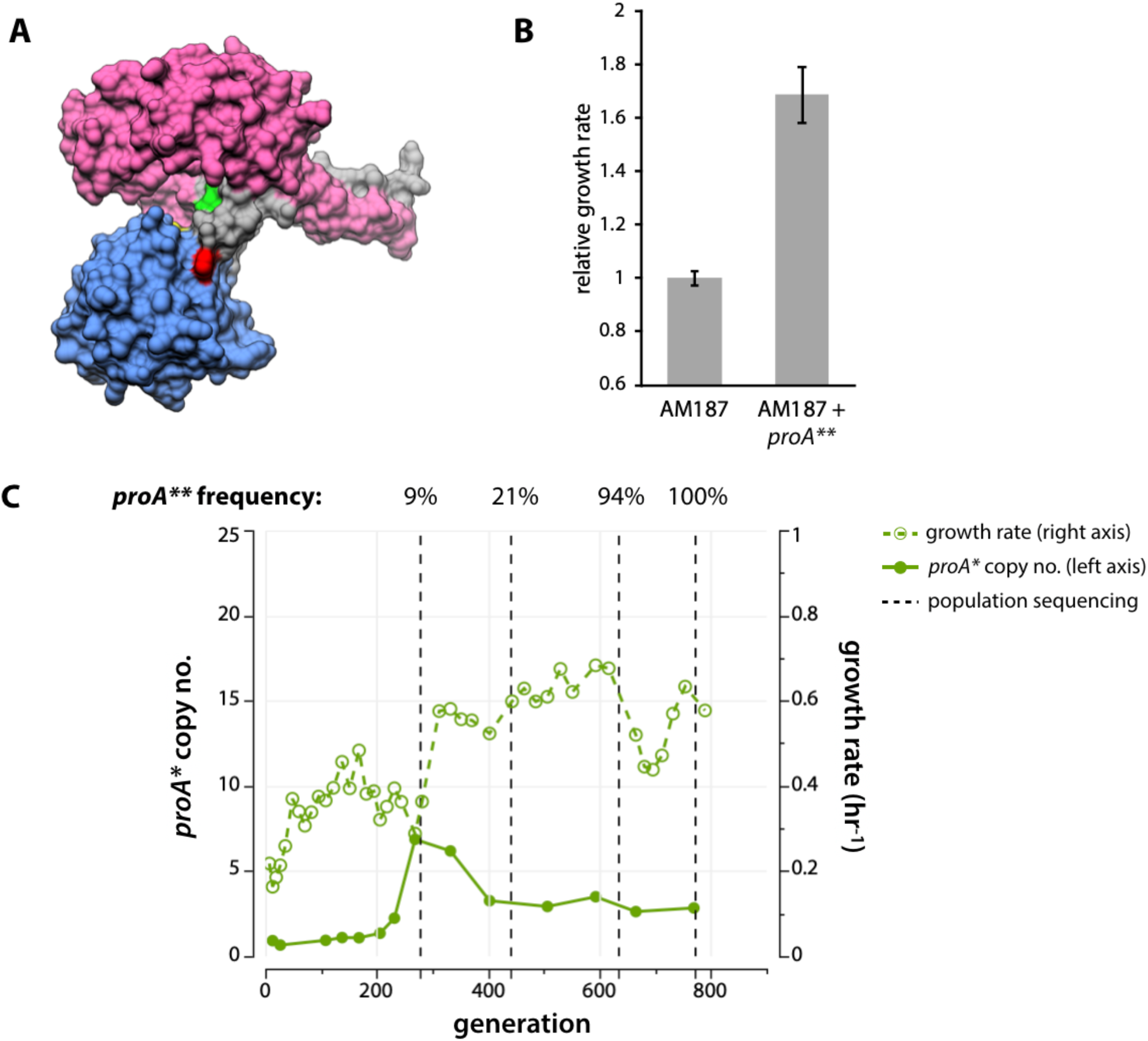
*proA\** acquired a beneficial mutation in population 3. (A) Crystal structure of *Thermotoga maritima* ProA (PDB 1O20) (Page et al., 2003). Yellow, catalytic cysteine; green, equivalent of *E. coli* ProA Glu383; red, equivalent of *E. coli* ProA Phe372; magenta, NADPH-binding domain; blue, catalytic domain; gray, oligomerization domain. (B) Change in growth rate when the mutation changing Phe372 to Leu is introduced into the genome of the parental strain (AM187). Error bars represent one standard error from the mean, N = 4. (C) Growth rate (right axis, dotted lines) and *proA\** copy number (left axis, solid lines) for population 3. Vertical dotted lines indicate when population genomic DNA was sequenced. The frequency of the *proA\*\** allele at each time point is noted above the plot.

The neo-ArgC and native ProA activities of wild-type, ProA*, and ProA** were assayed (in the reverse direction) with NAGSA and GSA, respectively (Table 2). The *k*_*cat*_/*K*_*M*_,_NAGSA_ for ProA** is 3.6-fold higher than that of ProA* and nearly 80-fold higher than that for ProA. In contrast, there is no difference between *k*_*cat*_/*K*_*M,GSA*_ for ProA* and ProA**.

**Table 2.** Kinetic parameters for GSA and NAGSA dehydrogenase activities of ProA, ProA*, and ProA**.

|  | GSA activity (ProA) |  |  | NAGSA activity (neo-ArgC) |  |  |
| --- | --- | --- | --- | --- | --- | --- |
| | $k_{cat}$<br>(s<sup>-1</sup>) | $K_M$<br>(mM) | $k_{cat}/K_{M,GSA}$<br>(M<sup>-1</sup> s<sup>-1</sup>) | $k_{cat}$<br>(s<sup>-1</sup>) | $K_M$<br>(mM) | $k_{cat}/K_{M,NAGSA}$<br>(M<sup>-1</sup> s<sup>-1</sup>) |
| **WT** | 16 ± 0.3 | 0.22 ± 0.01 | 72000 ± 2000 | 0.0083 ± 0.0009 | 0.30 ± 0.09 | 28 ± 9 |
| **ProA\***<br>**(E383A)** | 0.0076 ± 0.0008 | 0.20 ± 0.04 | 37 ± 8 | 0.046 ± 0.002 | 0.076 ± 0.009 | 610 ± 74 |
| **ProA\*\***<br>**(E383A**<br>**F372L)** | 0.023 ± 0.005 | 0.42 ± 0.14 | 55 ± 22 | 0.21 ± 0.01 | 0.095 ± 0.011 | 2200 ± 260 |

<sup>a</sup> Values reported were calculated from a nonlinear least squares regression of three replicates at each substrate concentration ± standard error.

To determine when the mutation that changes Phe372 to Leu in ProA* occurred, we performed sequencing of population genomic DNA at generations 270, 440, 630 and at the end of the adaptation (Figure 5C, 130×, 122×, 70× and 81× sequencing depth, respectively). *proA\*\** was present in 9% of the sequencing reads of the population by generation 270. By the time deamplification of *proA\** had occurred at generation 440, the frequency of *proA\*\** had risen to 21% of sequencing reads. By the end of the adaptation, *proA\*\** was fixed in the population, yet 3 copies remained in the genome, suggesting that ProA** does not have sufficient neo-ArgC activity to be present at a single copy in the genome.

The fact that a mutation that improved the neo-ArgC activity of ProA* occurred in only one population was surprising considering that ProA* is the weak-link enzyme limiting growth rate. Because the growth rates of all 8 populations improved substantially (Figure 3), mutations outside of the *proBA\** operon must also be contributing to fitness.

### Mutations outside of *proA\** improved fitness

Population genome sequencing at the end of the experiment and at several intermediate timepoints from populations with longer adaptations (Figure S1) revealed that the final populations contained 13-178 mutations present at frequencies ≥5%, 3-5 mutations at frequencies ≥30%, and 1-4 mutations (not including amplification of *proA\**) that were fixed (100%). We found several mutations in the same genes in different populations, suggesting that these mutations confer a fitness advantage (Supplementary File 1).

The first mutation to appear in all populations was either an 82 bp deletion in the *rph* pseudogene directly upstream of *pyrE* or a C➝T mutation in the intergenic region between *rph* and *pyrE*. PyrE is required for *de novo* synthesis of pyrimidine nucleotides. Both of these mutations have arisen in other *E. coli* evolution experiments, and have been shown to restore a known PyrE deficiency in the BW25113 *E. coli* strain (Blank et al., 2014; Bonekamp et al., 1984; Conrad et al., 2009; Jensen, 1993; Knöppel et al., 2018). Thus, these mutations are general adaptations to growth in minimal medium and do not pertain to the selective pressures caused by the weak-link enzyme ProA*.

A second early mutation in four of the populations occurred in *ygcB*. This mutation changes Ala390 to Val in Cas3, a nuclease/helicase in the Type I CRISPR/Cas system in *E. coli* (Howard et al., 2011). We introduced this mutation into the genome of the parent AM187 and compared the growth rates of the mutant and AM187 (Figure S2). Surprisingly, we saw no significant change in growth rate. Since this mutation appeared about the same time as the mutations upstream of *pyrE*, we wondered whether the *ygcB* mutation might only improve growth rate in the context of restored *pyrE* expression. Thus, we also tested the growth rate of a strain with the Cas3 mutation and the 82 bp deletion upstream of *pyrE*. Again, we saw no significant change in relative growth rate (Figure S2). Thus, the *ygcB* mutation is most likely a neutral hitchhiker. The most likely explanation for its prevalence is that it was present in a clade of the parental population that later rose to a high frequency when an additional beneficial mutation was acquired by one of its members.

### Mutations upstream of *argB* increase ArgB abundance

All eight final populations contained mutations in the intergenic region upstream of *argB* and downstream of *kan^r^*. These mutations were fixed in two populations, and present at frequencies of 9-82% in the other populations (Figure 6A).

**Figure 6.**
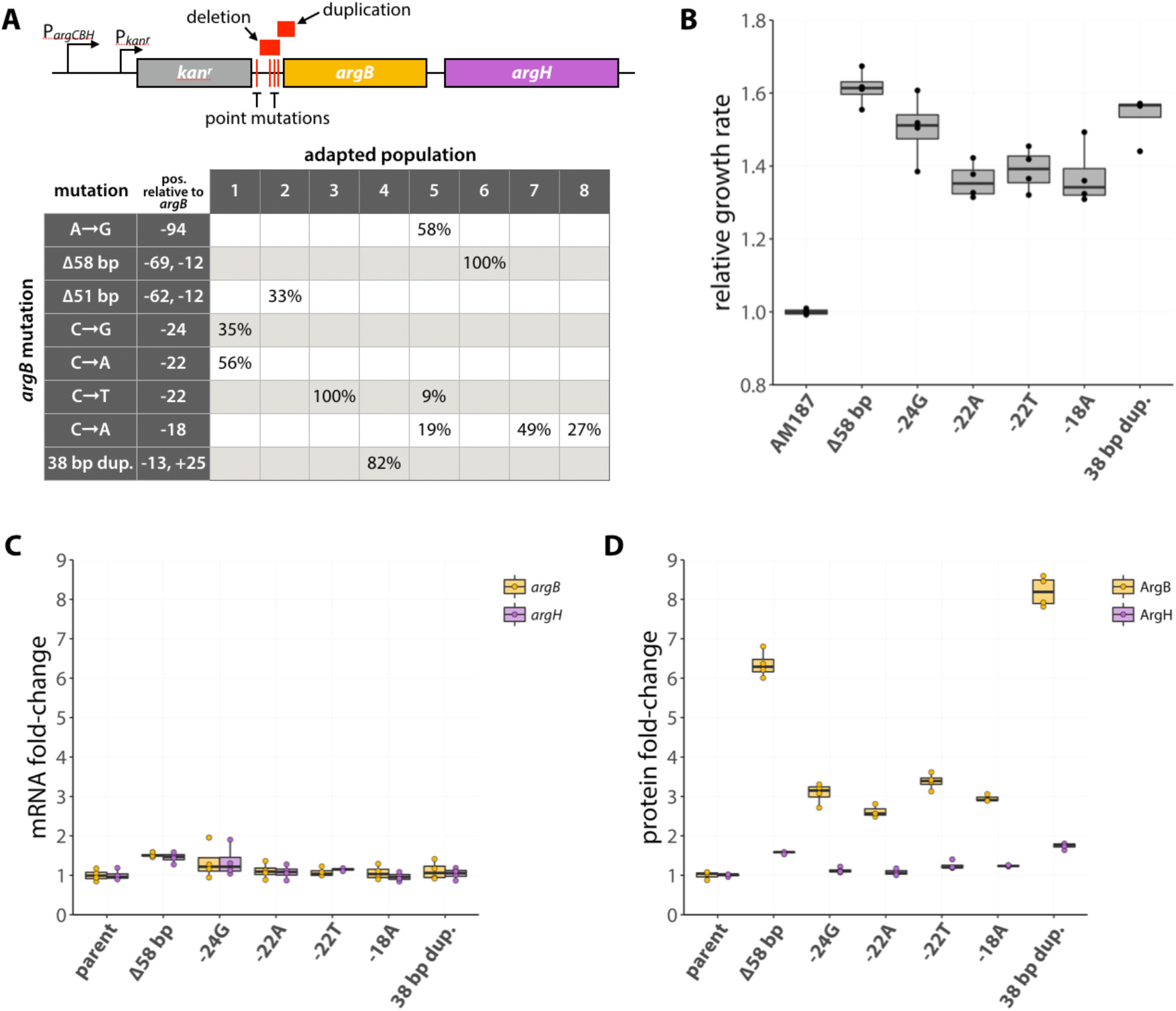
Several adaptive mutations occurred upstream of *argB*. (A) Locations of adaptive mutations (red) found upstream of *argB* (yellow) and *argH* (purple). *argC* was replaced with *kan*^*r*^ in the parental strain AM187, giving this operon two promoters, one native to the operon (P_*argCBH*_), and the other introduced with the *kan*^*r*^ gene (P_*kanr*_). The table shows the percentages of each adapted population that contained a given *argB* mutation at the final time point. Six of the *argB* mutations were introduced into the genome of the ancestral strain and changes in growth rate (B), gene expression (C), and protein abundance (D) were determined.

We reintroduced six of the mutations upstream of *argB* into the parental strain AM187. The mutations increased growth rate by 36-61% (Figure 6B). Levels of mRNAs for *argB* and *argH*, which is immediately downstream of *argB*, were little affected by the mutations (Figure 6C). However, levels of ArgB protein measured by label-free high resolution Orbitrap mass spectrometry increased 2.6-8.2-fold (Figure 6D). In contrast, ArgH levels increased only modestly. These data suggest that the mutations upstream of *argB* increase translational efficiency of *argB* mRNA.

Increased translational efficiency can be due to increased accessibility of the region surrounding the Shine-Dalgarno site and the start codon, which might be achieved in three ways: 1) a decrease in the stability of the lowest-free-energy secondary structure in this region (Bentele et al., 2013; Espah Borujeni et al., 2014; Goodman et al., 2013); 2) a change in the ensemble of mRNA structures such that the Shine-Dalgarno sequence is more likely to be single-stranded in multiple accessible structures (Espah Borujeni et al., 2014; Salis et al., 2009); or 3) a decrease in the folding rate of the mRNA such that a ribosome can bind to the unfolded mRNA emerging behind a preceding ribosome before the mRNA folds and obscures the Shine-Dalgarno sequence (a process called ribosomal drafting, (Espah Borujeni & Salis, 2016)). The *argB* mRNA, like 16% of γ-proteobacterial mRNAs (Scharff et al., 2011), lacks a canonical Shine-Dalgarno sequence, but the ribosome is expected to bind to a region encompassing the start codon and at least the upstream 8-10 nucleotides. We calculated the minimum free energy secondary structures of the 140-nucleotide RNA sequences encompassing the intergenic region affected by the various mutations through 33 bp downstream of the *argB* start codon (Figure S3). Direct comparisons of the predicted free energies of these structures are not particularly informative, especially for mutants with large deletions or duplications. However, for 5 of the 8 of the mutant structures, the probability that the 5’-UTR from about 20 nt upstream of the start codon is sequestered in a stem-loop in the lowest free-energy structure is decreased relative to the parent (Figure S3B); the increased accessibility of this region should increase translation efficiency. However, for three cases (−94 A➝G, −22 C➝A, and −18 C➝A), this region is equally or more likely to be sequestered in a stem-loop. The thermodynamic stability of this region is clearly not the only factor responsible for the effects of the mutations upstream of *argB*.

We also considered the possibility that mutations upstream of *argB* might increase expression by increasing ribosome drafting. Figure S4 shows the predicted folding times of RNA sequences encompassing 30 nucleotides downstream to 30 nucleotides upstream of the *argB* start codon (totaling 63 nucleotides) for each mutant except the −94 A→G mutant. Folding is predicted to be significantly slower for three of the seven mutant RNAs, which should increase the efficiency of translation. For four of the mutants, however, folding rate is similar to or even faster than that of the parental mRNA. For three of these mutants (the 58 bp deletion, −22 C→T, and the 38 bp duplication), the secondary structure prediction shown in Figure S3 suggests that the translation initiation region is less likely to be sequestered in a hairpin. Thus, the effects of 6 of the 8 mutations upstream of *argB* can be explained by a decrease in secondary structure stability around the ribosome binding site or a decrease in the folding rate of the mRNA in this region, or both. The effects of the −94 A➝G and −22 C➝A mutations, however, cannot be explained by either of these mechanisms.

### Mutations in *carB* either increase activity or impact allosteric regulation

We found eight different mutations in *carB* in six of the evolved populations: four missense mutations, three deletions (≥12 bp), and one 21 bp duplication (Figure 7A). CarB, the large subunit of carbamoyl phosphate synthetase (CPS), forms a complex with CarA to catalyze production of carbamoyl phosphate from glutamine, bicarbonate, and two molecules of ATP (Eq. [1]) (Rubino et al., 1986, 1987).

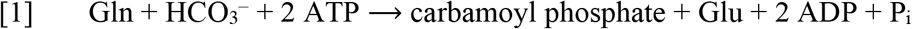

**Figure 7.**
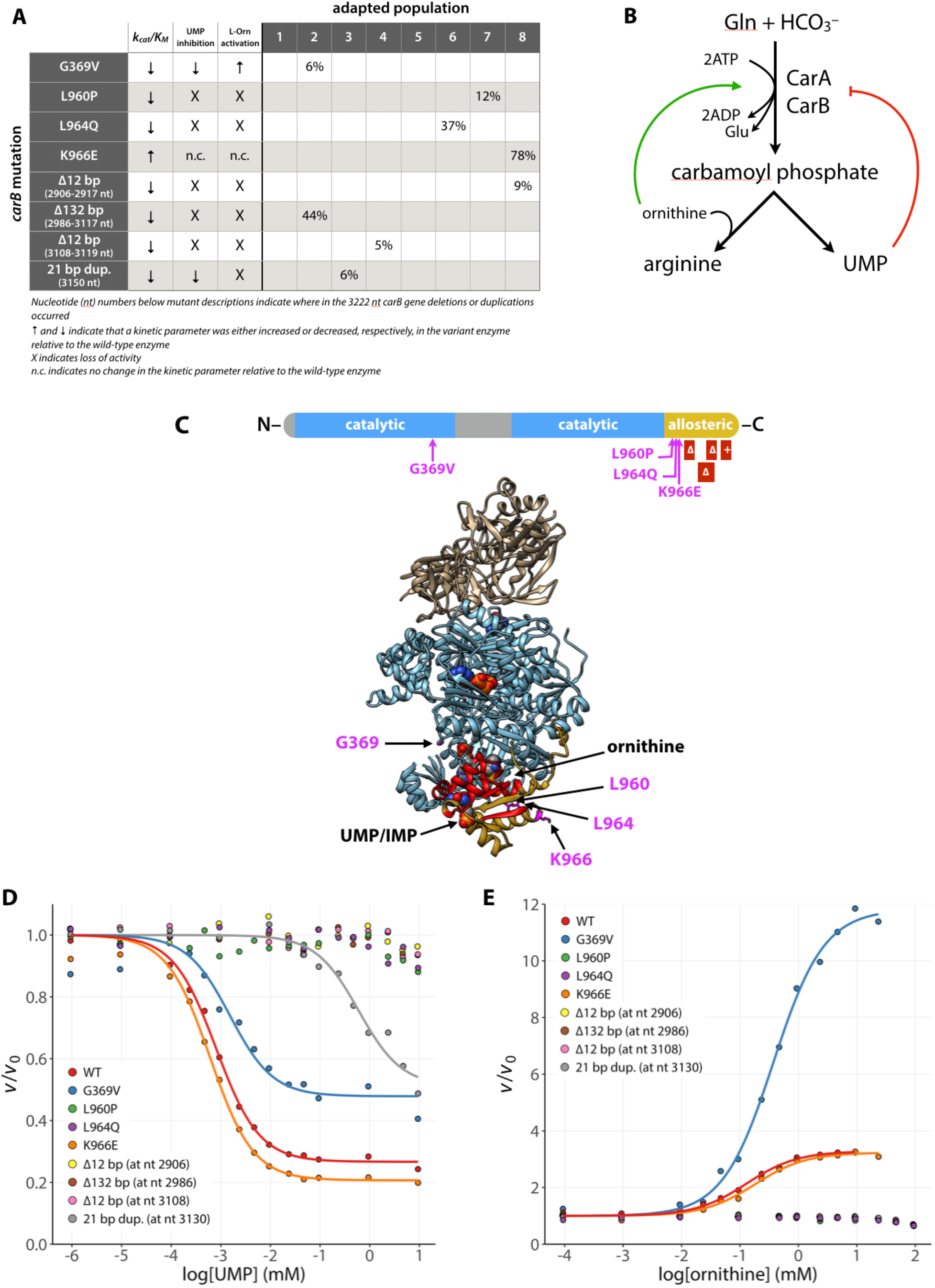
Several adaptive mutations occurred in *carB*. (A) Maximum percentage of each *carB* mutation found in the population at any time during the adaptation. (B) Allosteric regulation of carbamoyl phosphate synthetase. CarA and CarB are the small and large subunits of carbamoyl phosphate synthetase, respectively. (C) CarB functional domains (top) and crystal structure of *E. coli* CarAB (PDB 1CE8, bottom) (Thoden et al., 1999). Beige, CarA; blue, CarB; yellow, allosteric domain of CarB; red, residues that are deleted or duplicated in the adapted strains; magenta, point mutations that occur in the adapted strains. IMP and ornithine bound to the allosteric domain are shown as spheres. One of the two bound ATP molecules can be seen as spheres in the center of CarB. (D-E) Influence of UMP or L-ornithine concentration on the glutamine-dependent ATPase activity of CarAB mutants. Reaction rates are shown relative to *v*_0_, the reaction rate in the absence of ligand.

Synthesis of carbamoyl phosphate involves four reactions that take place in three separate active sites connected by a molecular tunnel of ~100 Å in length (Thoden et al., 2002). CarA catalyzes hydrolysis of glutamine to glutamate and ammonia (Eq. [2]), while CarB phosphorylates bicarbonate to form carboxyphosphate in its first active site (Eq. [3]). Ammonia from the CarA active site is channeled to CarB, where it reacts with carboxyphosphate to form carbamate (Eq. [4]). Carbamate migrates to a second active site within CarB where it reacts with ATP to form carbamoyl phosphate and ADP (Eq. [5]).

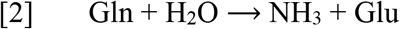

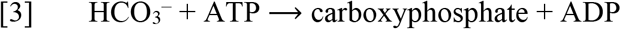

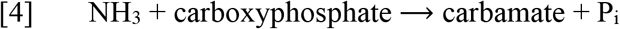

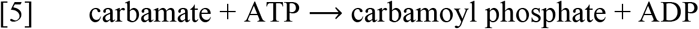

Carbamoyl phosphate feeds into both the pyrimidine and arginine synthesis pathways and its production is regulated in response to intermediates or products of both pathways (Figure 7B), as well as by IMP (a purine) (Pierrat & Raushel, 2002). It is inhibited by UMP (a pyrimidine) and moderately activated by IMP. UMP and IMP compete to bind the same region of CarB (Eroglu & Powers-Lee, 2002). The allosteric effects of UMP and IMP are dominated, however, by activation by ornithine. Ornithine, an intermediate in arginine synthesis that reacts with carbamoyl phosphate, binds to and activates CarB even when UMP is bound (Figure 7C) (Braxton et al., 1999; Eroglu & Powers-Lee, 2002).

Seven of the eight mutations found in *carB* affect residues in the allosteric domain of CarB. The lone mutation outside of the allosteric domain changes Gly369, which is immediately adjacent to the allosteric region in the 3D structure, to Val (Figure 7C).

The kinetic parameters for CPS activity [determined as the glutamine- and bicarbonate-dependent ATPase activity (Eq. [1])] of all eight CPS mutants are shown in Table 3. All mutations decreased *k*_*cat*_/*K*_*m,ATP*_ by 34-63%, with the exception of the mutation that changes Lys966 to Glu, which nearly doubles *k*_*cat*_/*K*_*m,ATP*_. None of the mutations affected the enzyme’s ability to couple ATP hydrolysis with carbamoyl phosphate production (Figure S5).

**Table 3.** Kinetic parameters^*a*^ for the glutamine-dependent ATPase reaction of wild-type and variant carbamoyl phosphate synthetases.

| Enzyme | $K_{M,ATP}$<br>(mM) | $k_{cat}$<br>(s<sup>-1</sup>) | $k_{cat}/K_{M,ATP}$<br>(M<sup>-1</sup> s<sup>-1</sup>) | UMP $K_d$<br>(μM) | UMP $a$ | ornithine $K_d$<br>(μM) | ornithine $a$ |
| --- | --- | --- | --- | --- | --- | --- | --- |
| WT | 1.05 ± 0.08 | 13.5 ± 0.3 | 12.9 (± 1.0) × 10<sup>3</sup> | 0.81 ± 0.04 | 0.27 ± 0.01 | 130 ± 7 | 3.28 ± 0.03 |
| G369V | 3.31 ± 0.25 | 21.5 ± 0.7 | 6.51 (± 0.54) × 10<sup>3</sup> | 1.53 ± 0.29 | 0.48 ± 0.02 | 372 ± 20 | 11.8 ± 0.13 |
| L960P | 1.12 ± 0.05 | 9.10 ± 0.15 | 8.12 (± 0.41) × 10<sup>3</sup> | na | na | na | na |
| L964Q | 1.25 ± 0.08 | 8.04 ± 0.17 | 6.41 (± 0.42) × 10<sup>3</sup> | na | na | na | na |
| K966E | 0.97 ± 0.06 | 20.6 ± 0.4 | 21.2 (± 1.4) × 10<sup>3</sup> | 0.61 ± 0.04 | 0.21 ± 0.01 | 181 ± 34 | 3.23 ± 0.08 |
| Δ12 bp (at nt 2906)<sup>b</sup> | 1.09 ± 0.06 | 4.80 ± 0.09 | 4.39 (± 0.25) × 10<sup>3</sup> | na | na | na | na |
| Δ132 bp (at nt 2986) | 1.16 ± 0.06 | 6.47 ± 0.11 | 5.57 (± 0.31) × 10<sup>3</sup> | na | na | na | na |
| Δ12 bp (at nt 3108) | 1.30 ± 0.10 | 5.86 ± 0.16 | 4.51 (± 0.37) × 10<sup>3</sup> | na | na | na | na |
| 21 bp dup. (at nt 3130) | 1.40 ± 0.12 | 9.70 ± 0.30 | 6.94 (± 0.64) × 10<sup>3</sup> | 597 ± 133 | 0.51 ± 0.03 | na | na |

<sup>a</sup> Values reported were calculated from a nonlinear least squares regression of data for three replicates at each substrate concentration ± standard error. Values for $K_d$ and $a$ for UMP and ornithine were determined by fitting the data to the following equation: $v/v_0 = (aL + K_d)/(L + K_d)$ , where L is the ligand concentration, v is the initial reaction
rate, $v_0$ is the initial reaction rate in the absence of ligand, $a$ is $v/v_0$ at infinite L, and $K_d$ is the apparent dissociation constant. No $K_d$ or $a$ values are given (indicated by *na*) when the inhibition by the allosteric ligand was too weak to measure.
<sup>b</sup> Nucleotide (nt) numbers refer to the position of deletions or duplications in *carB*.

We also measured the effect of mutations on UMP inhibition and ornithine activation of CPS (Table 3, Figure 7D-E). Regulation of the K966E variant, the enzyme for which *k*_*cat*_/*K*_*m,ATP*_ was nearly doubled, was minimally affected. Five of the variants showed a complete loss of allosteric regulation by both UMP and ornithine. The variant with the 21 bp duplication retained modest inhibition by UMP (~50% reduction of turnover at high UMP concentrations), but only at very high concentrations of UMP; the apparent *K*_*d,UMP*_ was increased by 740-fold. Similarly, G369V CPS retained partial inhibition by UMP (~50% reduction of turnover at high UMP concentrations). The apparent *K*_*d,UMP*_ of the G369V enzyme was only doubled, but this variant showed a 3.5-fold increase in activation at high ornithine concentrations.

## Discussion

Recruitment of promiscuous enzymes to serve new functions followed by mutations that improve the promiscuous activity has been a dominant force in the diversification of metabolic networks (Copley, 2017; Glasner et al., 2006; Khersonsky & Tawfik, 2010; O’Brien & Herschlag, 1999; Rauwerdink et al., 2016). New enzymes may be important for fitness or even survival when an organism is exposed to a novel toxin or source of carbon or energy, or when synthesis of a natural product can enable manipulation of competing organisms. This process also contributes to non-orthologous gene replacement, which can occur when a gene is lost during a time in which it is not required, but its function later becomes important again and is replaced by recruitment of a non-orthologous promiscuous enzyme (Albalat & Cañestro, 2016; Ferla et al., 2017; Juárez-Vázquez et al., 2017; Olson, 1999).

We have modeled a situation in which a new enzyme is required by deleting *argC*, which is essential for synthesis of arginine in *E. coli*. Previous work showed that the most readily available source of neo-ArgC activity that enables *∆argC E. coli* to grow on glucose as a sole carbon source is a promiscuous activity of ProA. However, a point mutation that changes Glu383 to Ala is required to elevate the promiscuous activity to a physiologically useful level. This mutation substantially damages the native function of the enzyme, creating an inefficient bifunctional enzyme whose poor catalytic abilities limit growth rate on glucose. It is important to note that the decrease in the efficiency of the native reaction may be a critical factor in the recruitment of ProA to serve as a neo-ArgC because it will diminish inhibition of the newly important reaction by the native substrate (Khanal et al., 2015; McLoughlin & Copley, 2008).

When we previously carried out short-term adaptation of *∆argC proA* E. coli* in glucose, we observed four mutations that increased fitness by increasing expression of *proA\**: a G➝A substitution in the −35 promoter region (M1) and a C➝T substitution in the −10 promoter region (M2) upstream of the *proBA\** operon, a synonymous mutation in *proB* that strengthens a cryptic promoter upstream of *proA\** (M3), and amplification of a region of the genome that includes the *proBA\** operon (Kershner et al., 2016). The point mutations occurred independently, never together. Amplification occurred even after acquisition of M1, M2, or M3. The amplification of *proA\** suggests that it has the potential to diverge to restore efficient ProA function and evolve an efficient neo-ArgC.

We chose to carry out longer-term evolution of a *∆argC proA\** strain in glucose in the presence of proline to specifically address the evolution of an efficient neo-ArgC. We introduced the M2 promoter mutation in the parent strain to ensure that all populations had the same promoter mutation during adaptation. After 470-1000 generations of adaptation, growth rate had increased ~3-fold in all eight replicate cultures. We have focused on five types of genetic changes that clearly increase fitness: (1) mutations upstream of *pyrE*; (2) amplification of a variable region of the genome surrounding the *proBA\** operon; (3) a mutation in *proA\** that changes Phe372 to Leu; (4) mutations upstream of *argB*; and (5) mutations in *carB*. (Each of the final populations contains additional mutations that may also contribute to fitness, but these mutations were typically found in low abundance and/or in only one population.) The *pyrE* mutations have previously been shown to be a general adaptation of *E. coli* BW25113 to growth in minimal medium (Blank et al., 2014; Conrad et al., 2009; Jensen, 1993; Knöppel et al., 2018). The other four types of mutations are specific adaptations to the bottleneck in arginine synthesis caused by substitution of the weak-link enzyme ProA* for ArgC. Interestingly, only two of these – gene amplification and the mutation in *proA\** – directly involved the weak-link enzyme ProA*.

Surprisingly, we saw evolution of *proA\** towards a more efficient neo-ArgC in only one population (Table 2). In this population, *proA* copy number dropped from ~7 to ~3 within 100 generations (Figure 5C). This pattern is consistent with the IAD model; copy number is expected to decrease as mutations increase the efficiency of the weak-link activity. However, the fact that copy number did not return to one implies that the neo-ArgC activity of ProA** is not sufficient to justify a single copy of the gene.

Because ~3 copies of *proA\*\** remained in the population (Figure 5C) and the progenitor *proA\** was not detectable, all copies in the amplified array have clearly acquired the mutation that changes Phe372 to Leu – i.e. the more beneficial *proA\*\** allele has “swept” the amplified array. This observation has important implications for the IAD model. In the original conception of the IAD model, it was proposed that amplification of a gene increases the opportunity for different beneficial mutations to occur in different alleles, and then for recombination to shuffle these mutations (Bergthorsson et al., 2007; Francino, 2005). Both phenomena would increase the rate at which sequence space can be searched and thereby increase the rate at which a new enzyme evolves. In order for this to occur, however, it would be necessary for individual alleles to acquire different beneficial mutations before recombination occurred. This scenario is inconsistent with the relative frequencies of point mutations and recombination between large homologous regions in an amplified array (Anderson & Roth, 1981; Reams et al., 2010). Point mutations occur at a frequency between 10^−9^ and 10^−10^ per nucleotide per cell division depending on the genomic location (Jee et al., 2016), and thus between 10^−6^ and 10^−7^ per gene per cell division for a gene the size of *proA*. If 10 copies of an evolving gene were present, then the frequency of mutation in a single allele would be between 10^−5^ and 10^−6^ per cell division. Homologous recombination after an initial duplication event is orders of magnitude more frequent, occurring in ~1 of every 100 cell divisions (Reams et al., 2010). Thus, homologous recombination between replicating chromosomes in a cell could result in a selective allelic sweep long before a second beneficial mutation occurs in a different allele in the amplified array. Figure 8 depicts how this could occur after acquisition of an advantageous mutation in one allele in an amplified array. This is indeed the result that we observed; heterozygosity among *proA\** alleles was lost within 500 generations (Figure 5C). More recent papers depict selective amplification of beneficial alleles before acquisition of additional mutations (Andersson et al., 2015; Näsvall et al., 2012); our results support this revision of the original IAD model. It is possible that alleles encoding enzymes that are diverging toward two specialists might recombine to explore combinations of mutations. However, such recombination might not accelerate evolution, as mutations that lead toward one specialist enzyme would likely be incompatible with those that lead toward the other specialist enzyme.

**Figure 8.**
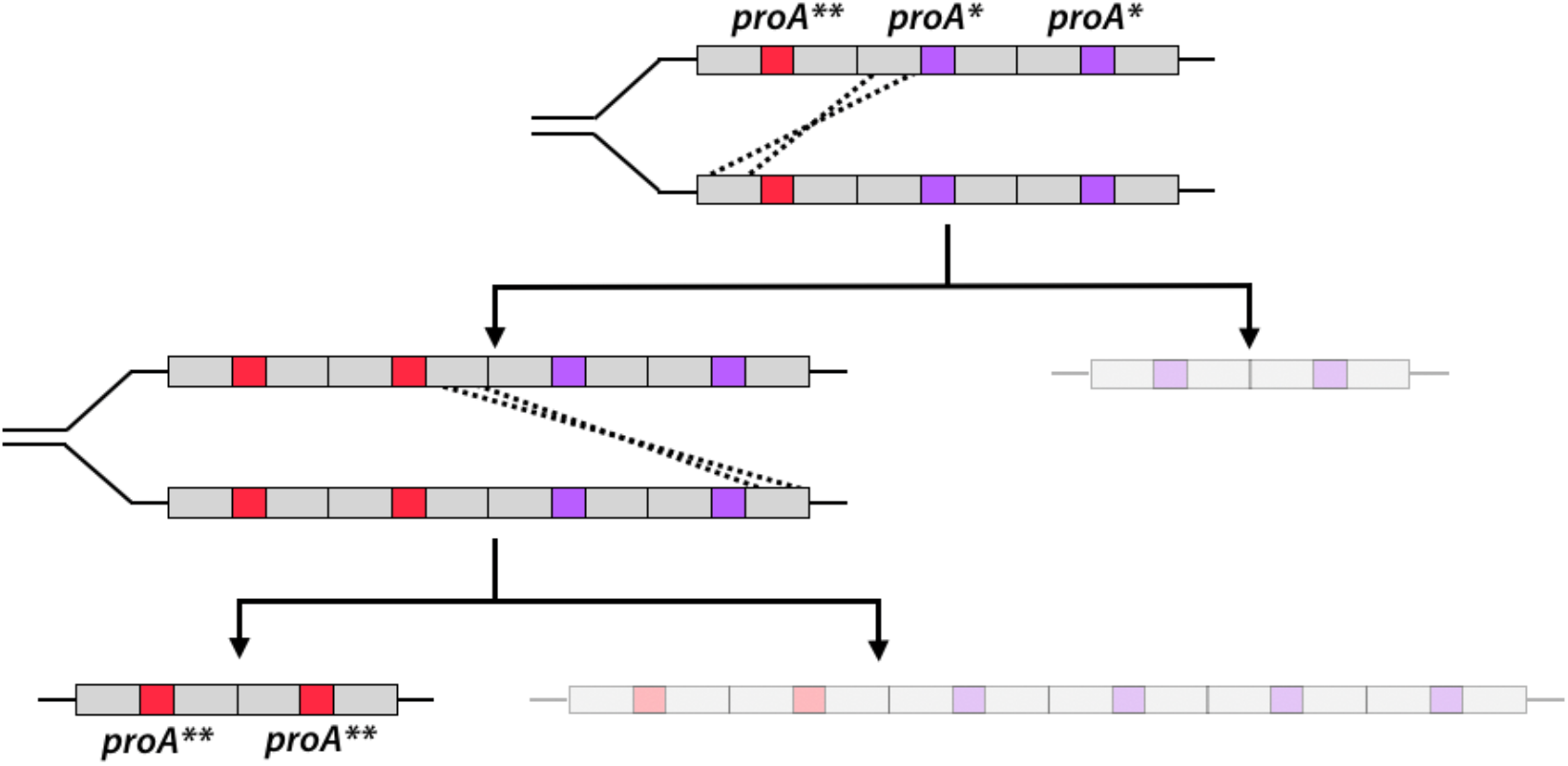
Homologous recombination of an amplified *proA\** array with one *proA\*\** allele can rapidly lead to a daughter cell with only *proA\*\** alleles. Each arrow represents one cell duplication event and shows the genotypes of the two resulting daughter cells. The less-fit daughter cell from each recombination event is grayed out.

While growth rate improved substantially in all populations, a beneficial mutation in *proA\** arose in only one, suggesting that either mutations that improve the neo-ArgC activity are uncommon, or that their fitness effects are smaller than those caused by mutations elsewhere in the genome that also improve arginine synthesis. We identified two primary mechanisms that apparently improve arginine synthesis without affecting the efficiency of the weak-link enzyme ProA* itself.

ArgB (*N*-acetylglutamate kinase) catalyzes the second step in arginine synthesis by phosphorylating *N*-acetylglutamate to form NAGP, the native substrate for ArgC and a secondary substrate for ProA* (Figure 2). We identified eight mutations upstream of *argB*; the six we tested improved growth rate by 36-61% and increased the abundance of ArgB by 2.6-8.2-fold. Notably, ArgB levels were increased even though the levels of *argB* mRNA were unchanged (Figure 6). The increase in protein levels without a concomitant increase in mRNA levels suggests that these mutations impact the efficiency of translation. Secondary structure around the translation initiation site plays a key role because this region must be unfolded in order to bind to the small subunit of the ribosome (Hall et al., 1982; Scharff et al., 2011). Indeed, a study of the predicted secondary structures of 5000 genes from bacteria, mitochondria and plastids, many of which lack canonical Shine-Dalgarno sequences (as does *argB*), showed that secondary structure around the start codon is markedly less stable than up- or down-stream regions (Bentele et al., 2013; Espah Borujeni et al., 2014; Goodman et al., 2013; Scharff et al., 2011). Our computational studies of the effect of mutations on the predicted lowest free energy secondary structures of the region surrounding the start codon of *argB* suggest that the thermodynamic stability of this region plays a role in the beneficial effects of most of the observe mutations (Figure S3). In addition, some of the mutations slow the predicted rate of mRNA folding around the start codon, which would increase the probability of ribosomal drafting (Figure S4). Both effects would lead to an increase in ArgB abundance. An increase in the level of ArgB should increase the concentration of the NAGP substrate for the weak-link ProA*, thereby pushing material through this bottleneck in the arginine synthesis pathway.

The adaptive mutations in *carB* increase catalytic turnover, decrease inhibition by UMP, or increase activation by ornithine of CPS. All of these effects should increase the level of CPS activity in the cell and consequently the level of carbamoyl phosphate. Why would this be advantageous? Carbamoyl phosphate reacts with ornithine, which will be in short supply due to the upstream ProA* bottleneck, to generate citrulline (Legrain & Stalon, 1976) (Figure 9). If ornithine transcarbamoylase is not saturated with respect to carbamoyl phosphate, then increasing carbamoyl phosphate levels should increase citrulline production and thereby increase flux into the lower part of the arginine synthesis pathway. Although we do not know whether ornithine transcarbamoylase is saturated with respect to carbamoyl phosphate *in vivo* (the *K*_*M*_ for carbamoyl phosphate is 360 µM (Baur et al., 1990), but the intracellular concentration of carbamoyl phosphate is unknown), the occurrence of so many mutations that increase carbamoyl phosphate synthase activity supports the notion that they lead to an increase in carbamoyl phosphate that potentiates flux through the arginine synthesis pathway.

**Figure 9.**
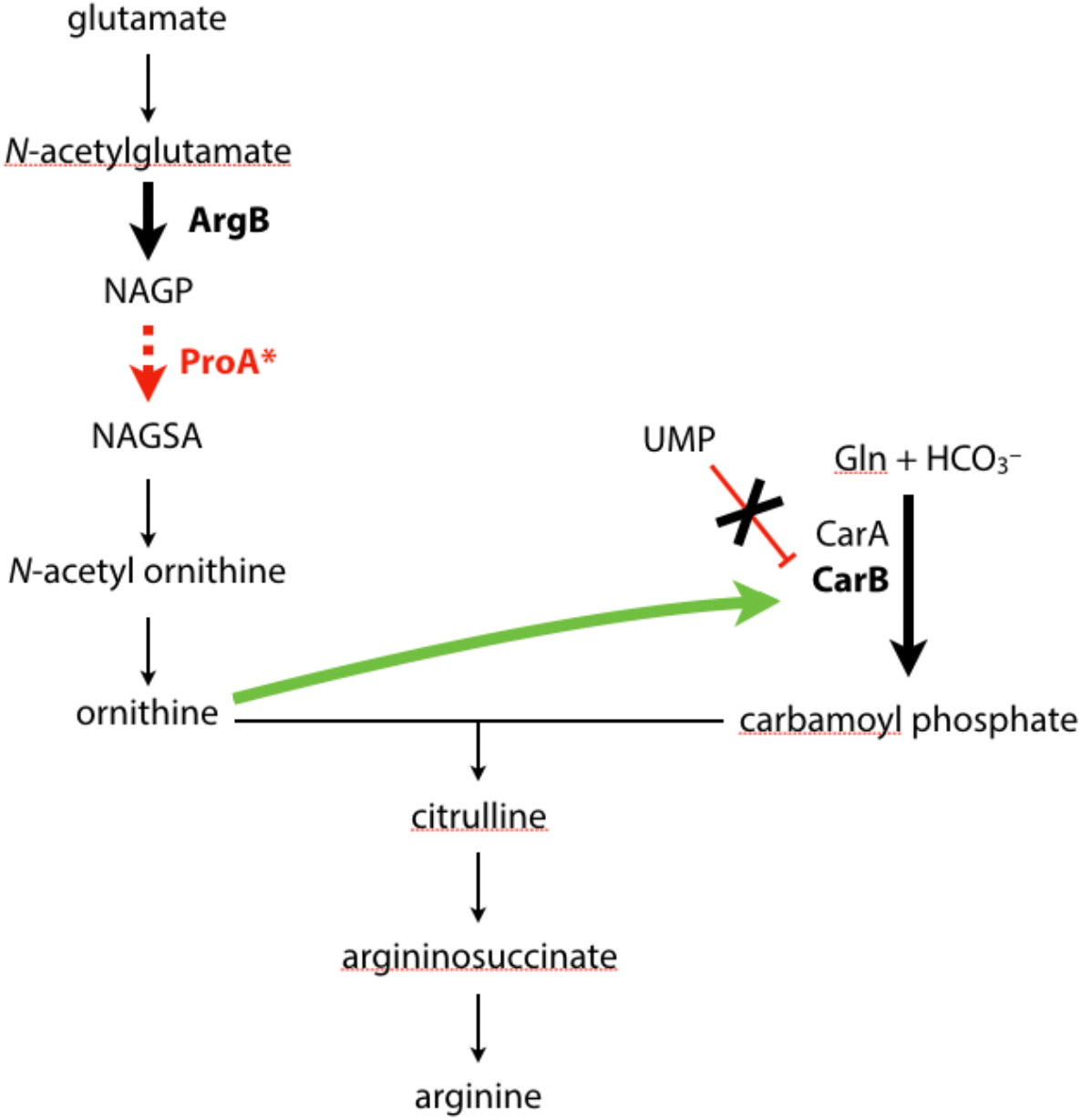
Adaptive mutations are predicted to increase flux through the arginine synthesis pathway. The pathway bottleneck and weak-link enzyme ProA* is shown in red. Steps in the arginine synthesis pathway that are affected by adaptive mutations are highlighted in bold type.

The majority of adaptive mutations we observed in *carB* cause loss of the exquisite allosteric regulation that controls flux through this important step in pyrimidine and arginine synthesis. This tight regulation likely evolved due to the energetically costly reaction catalyzed by CPS, which consumes two ATP molecules. While a constitutively active CPS is beneficial in the short term to improve arginine synthesis, it will likely be detrimental once arginine production no longer limits growth. We term mutations such as those observed in *carB* “expedient” mutations because they provide a quick fix when cells are under strong selective pressure, but at a cost to a previously well-evolved function. The damage caused by expedient mutations may be repairable later by reversion, compensatory mutations or horizontal gene transfer. Interestingly, the latter two repair processes may contribute to sequence divergence between organisms that has typically been attributed to neutral drift, but rather may be due to scars left from previous selective processes.

A particularly striking conclusion from this work is that most of the mutations that improved fitness under these strong selective conditions did not impact the gene encoding the weak-link enzyme, but rather adjusted fluxes in the metabolic network to compensate for the bottleneck in metabolism by other mechanisms. Not surprisingly, the process of evolution of a new enzyme by gene duplication and divergence does not take place in isolation, but is inextricably intertwined with mutations in the rest of the genome. The ultimate winner in a microbial population exposed to a novel selective pressure that requires evolution of a new enzyme may be the clone that succeeds in evolving an efficient enzyme that no longer limits fitness while accumulating the least damaging, or at least the most easily repaired, expedient mutations.

---

**Figure S1.**
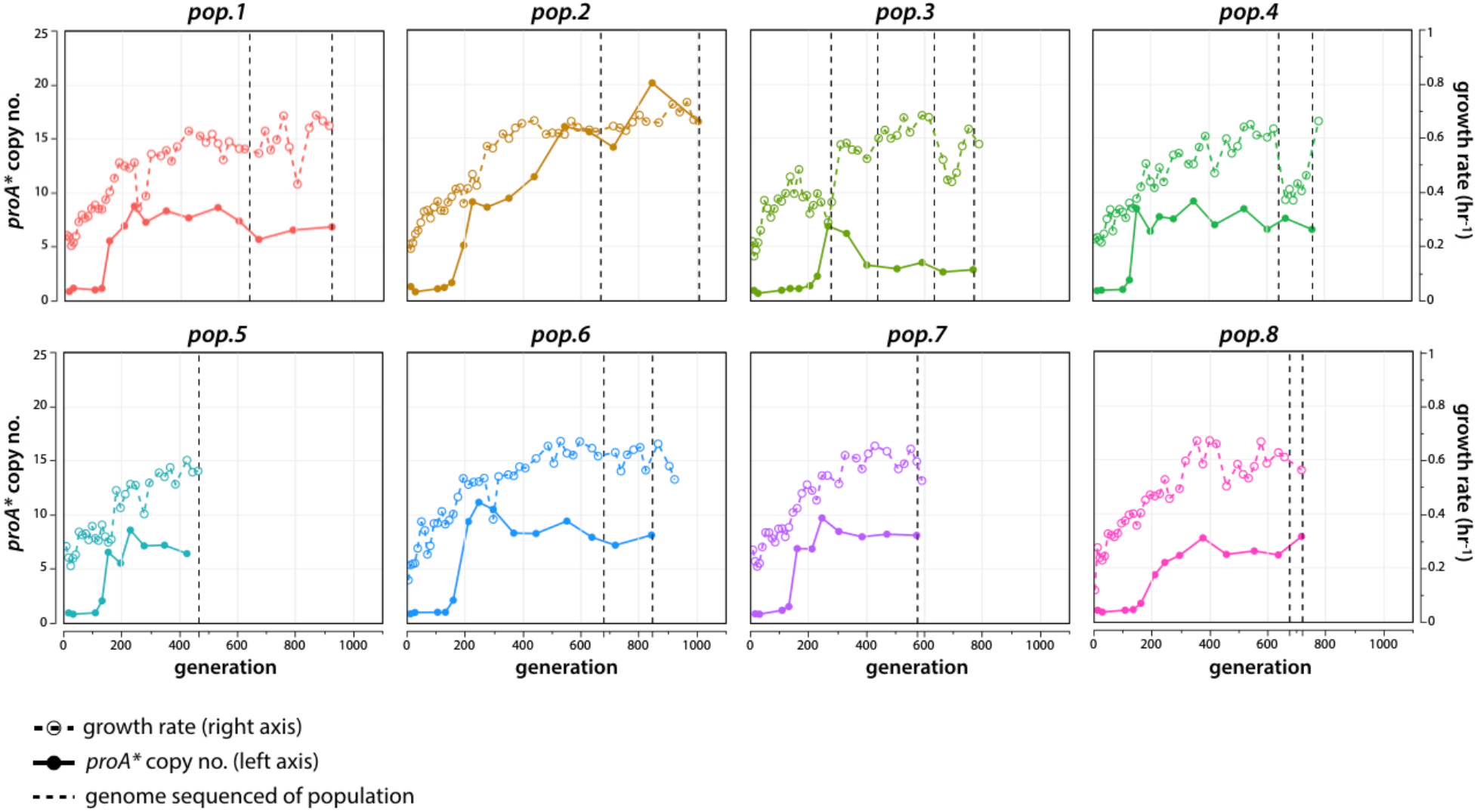
Growth rate (right axis, dotted lines) and *proA\** copy number (left axis, solid lines) for each adapted population. Vertical dotted lines indicate when population genomic DNA was sequenced.

**Figure S2.**
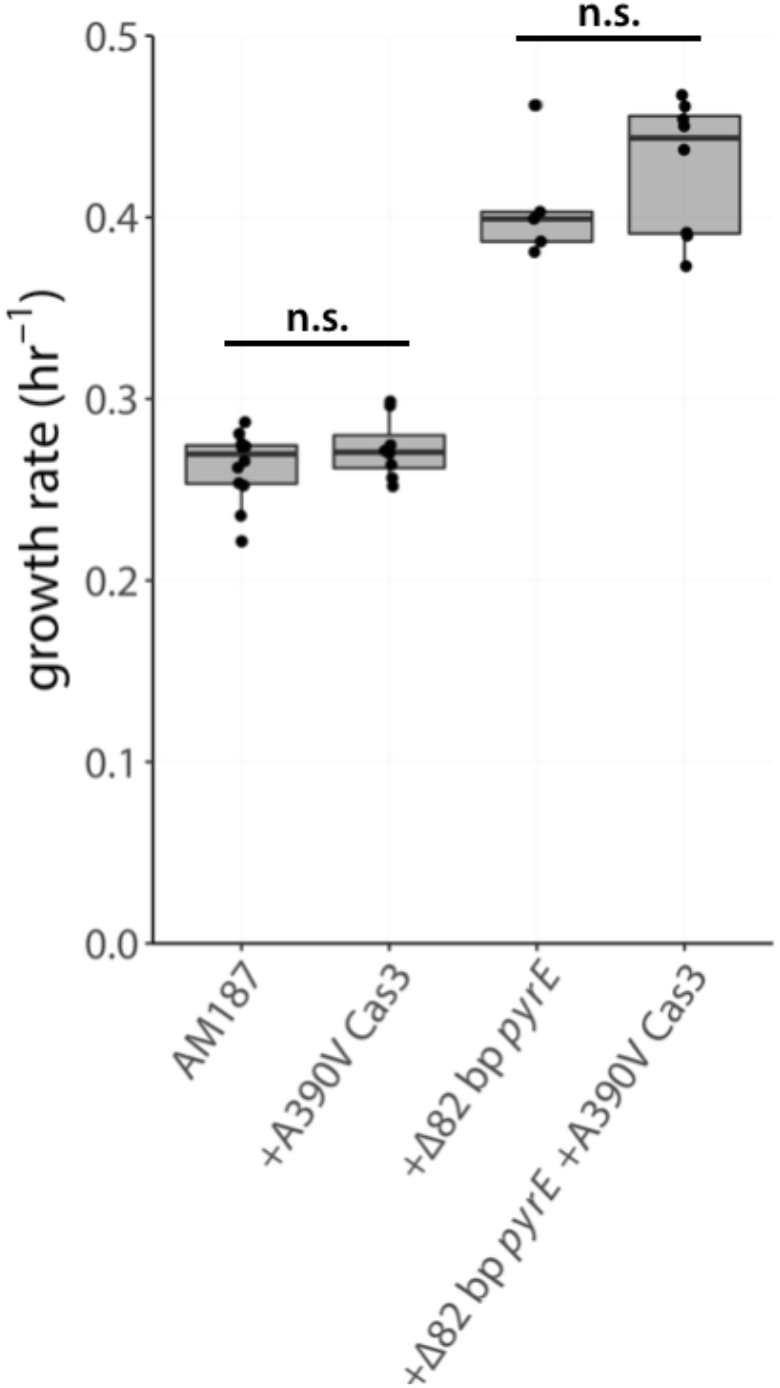
Effects of the A390V Cas3 and *rph-pyrE* mutations on growth rate of the parental strain AM187. (All *p* values are >0.1 by a one-tailed Student’s *t*-test).

**Figure S3.**
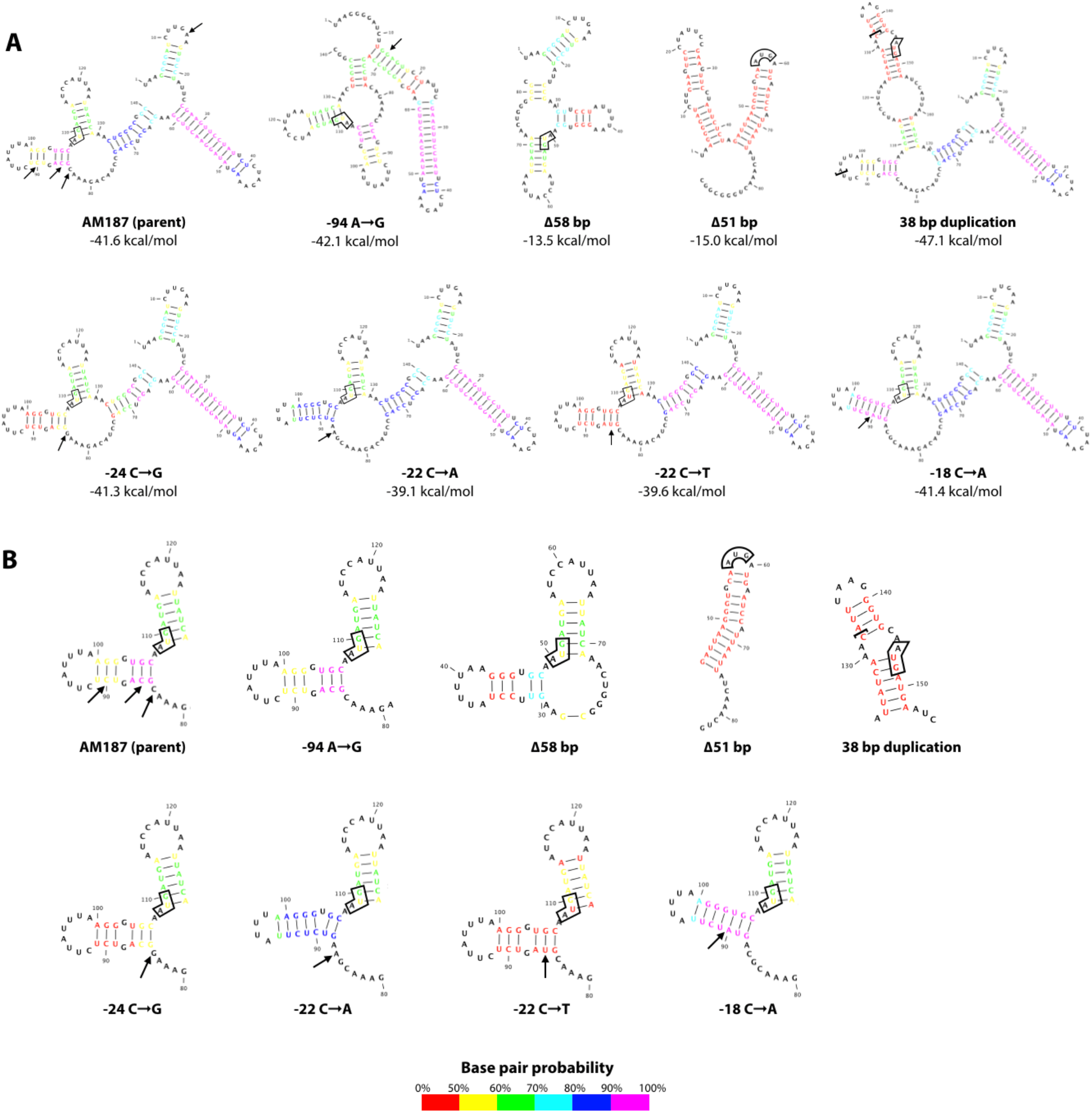
*argB* mRNA secondary structure comparison. (A) The entire intergenic region between *kan*^r^ and *argB* plus 33 nucleotides of the *argB* coding region was used for RNA structure prediction. The start codon for *argB* is boxed in each structure. Arrows in the AM187 structure indicate where an adaptive point mutation occurred. Arrows in mutant structures indicate which nucleotide was changed. In the 38 bp duplication mutant structure, the brackets contain the 38 inserted nucleotides. (B) Same as in (A), but showing only the region surrounding the *argB* start codon.

**Figure S4.**
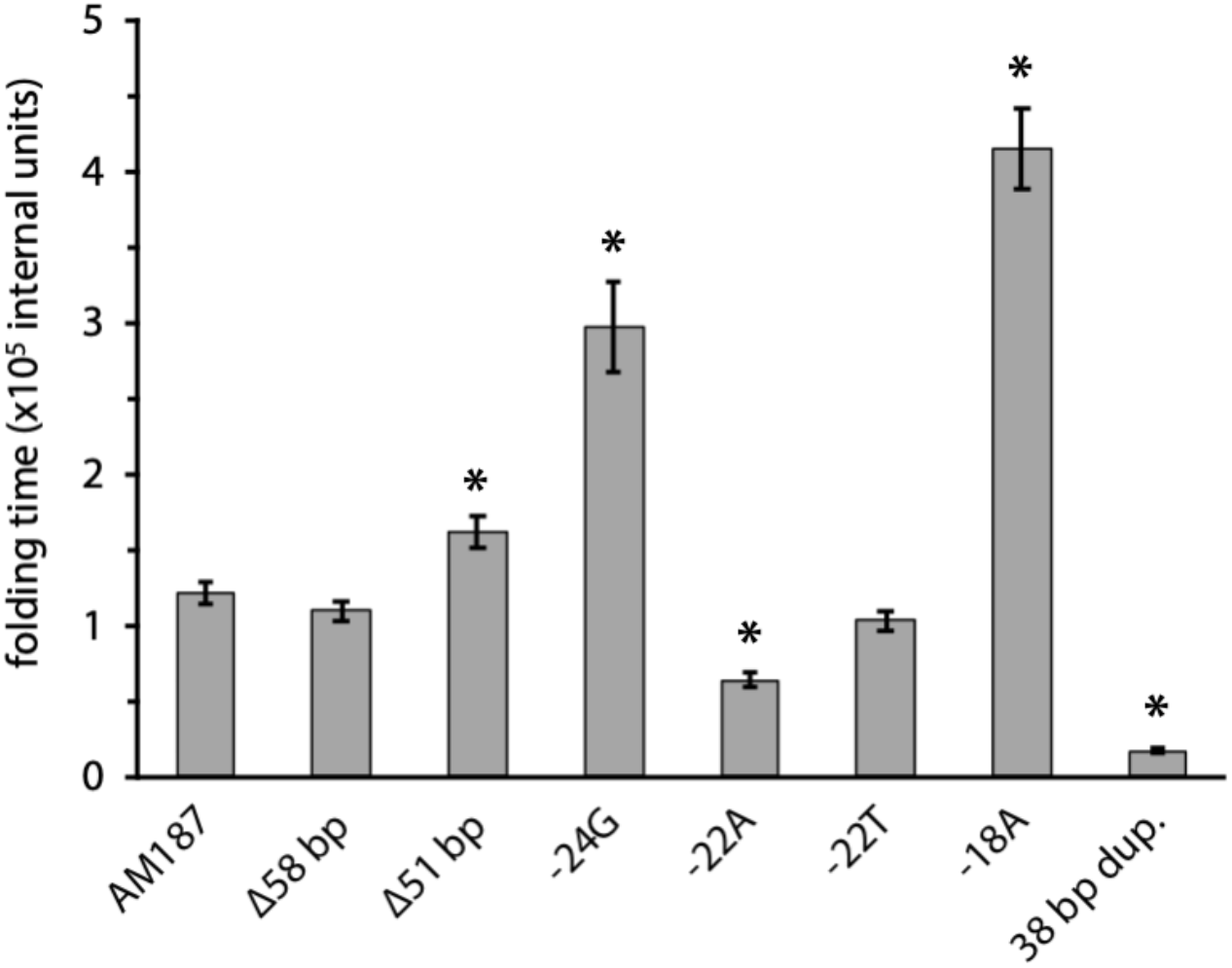
Folding time of a region encompassing 30 nucleotides upstream of the *argB* start codon plus 33 nucleotides of the *argB* coding region based on Kinfold simulations (Wolfinger et al., 2004). Error bars represent one standard error from the mean, N = 500.

**Figure S5.**
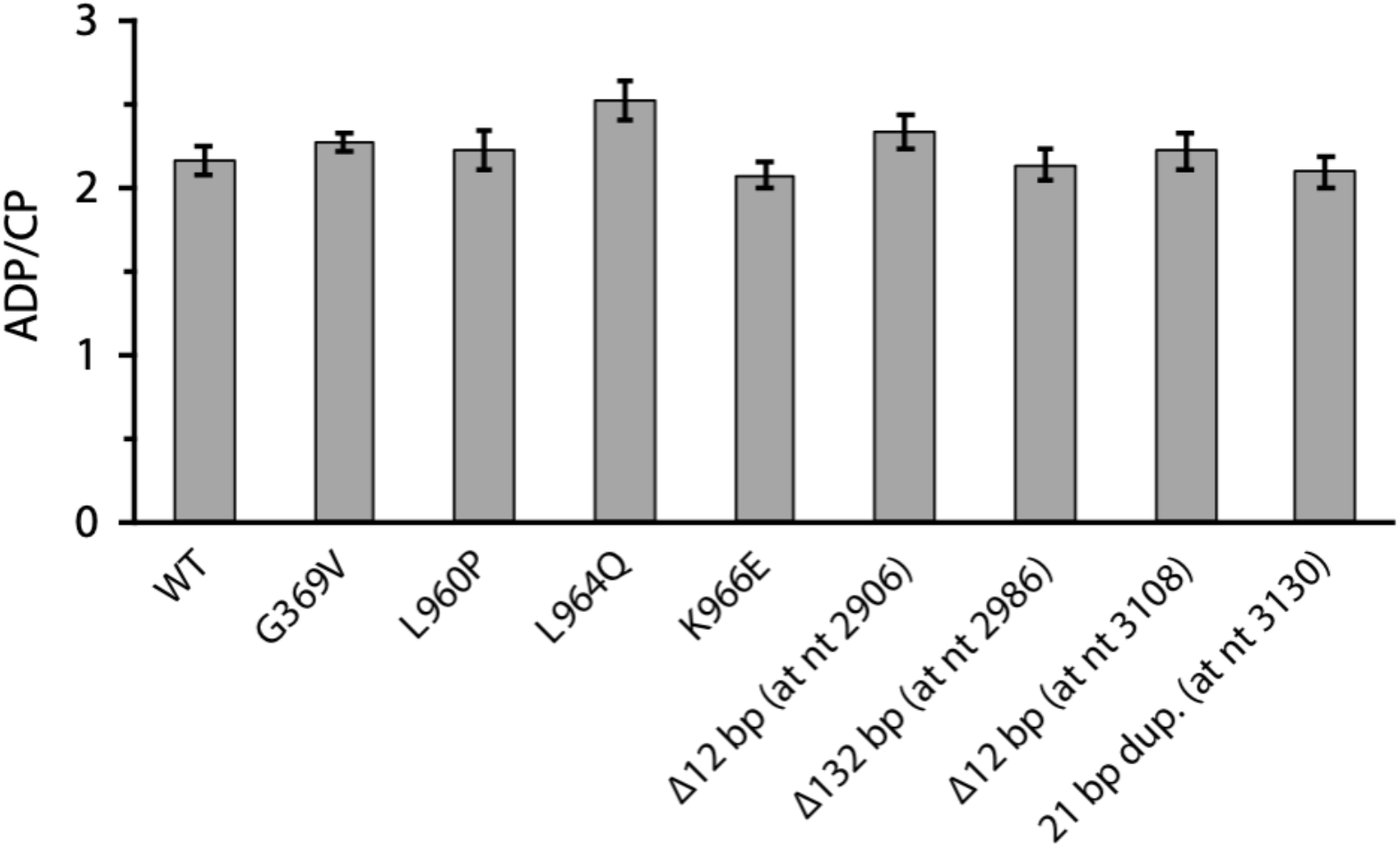
Ratio of ADP to carbamoyl phosphate (CP) produced by wild-type and mutant carbamoyl phosphate synthetases.

## Materials and Methods

### Materials

Common chemicals were purchased from Sigma-Aldrich (St. Louis, MO) and Fisher Scientific (Fair Lawn, NJ).

NAGSA was synthesized enzymatically from *N*-acetylornithine using *N*-acetylornithine aminotransferase (ArgD) in a 25 mL reaction as described previously by Khanal et al. 2015 and stored at −70 °C. NAGSA concentrations were determined using the *o*-aminobenzaldehyde assay as described previously (Albrecht et al., 1962; Mezl & Knox, 1976).

GSA was synthesized enzymatically from L-ornithine using *N*-acetylornithine aminotransferase (ArgD) as described previously by Khanal et al. 2015 and stored at −70 °C. GSA concentrations were determined using the *o*-aminobenzaldehyde assay as described previously (Albrecht et al., 1962; Mezl & Knox, 1976).

Plasmids and primers used in this study are listed in Tables S1 and S2, respectively.

### Strains and culture conditions

Strains used in this study are listed in Table 1. *E. coli* cultures were routinely grown in LB medium at 37 °C with 20 µg/mL kanamycin, 100 µg/mL ampicillin, 20 µg/mL chloramphenicol, or 10 µg/mL tetracycline, as required. Adaptation of strain AM187 was performed at 37 °C in M9 minimal medium containing 0.2% glucose, 0.4 mM proline, and 20 µg/mL kanamycin (Adaptation Medium).

### Strain construction

The parental strain for the adaptation experiment (AM187) was constructed from the Keio collection *argC*::*kan*^r^ *E. coli* BW25113 strain (Baba et al., 2006). The *fimAICDFGH* and *csgBAC* operons were deleted (to slow biofilm formation), and the M2 *proBA* promoter mutation (Kershner et al., 2016) and the point mutation in *proA* that changes Glu383 to Ala (McLoughlin & Copley, 2008) were inserted into the genome using the scarless genome editing technique described in Kim et al. 2014. We initially hoped to measure *proA\** copy number during adaptation using fluorescence, although ultimately qPCR proved to be a better approach. Thus, we inserted *yfp* downstream of *proA\** under control of the P3 promoter (Mutalik et al. 2013) and with a synthetically designed ribosome binding site (Espah Borujeni et al., 2014; Salis et al., 2009). A double transcription terminator (BioBrick Part: BBa_B0015) was inserted immediately downstream of *proBA\** to prevent read-through transcription of *yfp*. We also inserted a NotI cut site immediately downstream of *proA\** to enable cloning of individual *proA\** alleles after amplification if necessary. A Fis binding site located 32 bp downstream of *proA* was preserved because it might impact *proA* transcription. The NotI-2×Term-*yfp* cassette was inserted downstream of *proA\** using the scarless genome editing technique described in Kim et al. 2014. The genome of the resulting strain AM187 was sequenced to confirm that there were no unintended mutations and deposited to NCBI GenBank under accession number CP037857.

Most mutations observed during the adaptation experiment were introduced into the parental AM187 strain using the scarless genome editing protocol described in Kim et al. 2014. This protocol is preferable to Cas9 genome editing for introduction of point mutations and small indels because it does not require introduction of synonymous PAM mutations that have the potential to affect RNA structure. The 58 bp deletion upstream of *argB* and the 82 bp deletion in *rph* were introduced using Cas9-induced DNA cleavage and *λ* Red recombinase-mediated homology-directed repair with a linear DNA fragment. Sequences of the protospacers and mutations cassettes used for Cas9 genome editing procedures are listed in Tables S3 and S4. Details of the genome editing procedures are provided in the supplementary material.

### Laboratory adaptation

Adaptation of strain AM187 in Adaptation Medium was carried out in eight replicate tubes in a custom turbidostat constructed as described by Takahashi et al. 2015. To start the adaptation, strain AM187 was grown to exponential phase (OD_600_ = 0.7) in LB/kanamycin at 37 °C. Cells were centrifuged at 4,000 × *g* for 10 min at room temperature and resuspended in an equal volume of PBS. The suspended cells were washed twice more with PBS and resuspended in PBS. This suspension was used to inoculate all eight turbidostat chambers to give an initial OD_600_ of 0.01 in 14 mL of Adaptation Medium. The turbidostat was set to maintain an OD_650_ of 0.4 by diluting individual cultures with an appropriate amount of fresh medium every 60 seconds.

A 3 mL portion of each population was collected every 2-3 days; 800 µl was used to make a 10% glycerol stock, which was then stored at −70 °C. The remaining sample was pelleted for purification of genomic DNA using the Invitrogen PureLink Genomic DNA Mini Kit according to the manufacturer’s protocol.

The turbidostat occasionally malfunctioned, requiring the adaptation to be paused. When this occurred, the populations were subjected to centrifugation at 4,000 × *g* for 10 min at room temperature and the pelleted cells were resuspended in 1.6 mL of Adaptation Medium. Half of the resuspension was used to make a −70 °C 10% glycerol stock, and the other half to purify genomic DNA. When the turbidostat was fixed, the frozen stock was thawed and the cells were collected by centrifugation at 16,000 × *g* for 1 min at room temperature. The pelleted cells were resuspended in 1 mL of PBS, washed, and resuspended in 500 µL of Adaptation Medium. The entire resuspension was used to inoculate the appropriate chamber of the turbidostat. Sometimes the adaptation had to be restarted from a frozen stock of a normal sample (as opposed to the entire population as just described), resulting in a more significant population bottleneck. In this case, the entire frozen stock was thawed and only 700 µL washed as described above to be used for the inoculation. The remaining 300 µL of the glycerol stock were re-stored at −70 °C in case the frozen stock was needed for downstream analysis.

### Calculation of growth rate and generations during adaptation

The turbidostat takes an OD_650_ reading every ~3 sec and dilutes the cultures every 60 seconds. Thus, readings between dilutions can be used to calculate an average growth rate each day based on the following equation:

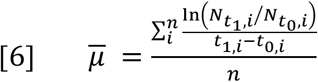

where 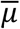 is the average growth rate in hr^−1^, *n* is the number of independent growth rate calculations within a given 24 hr period, 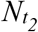 is the OD_650_ reading right before the dilution, 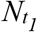is the OD_650_ reading right after the dilution, and *t*_1_ and *t*_0_ are the times at which the OD_650_ was measured. The number of generations per day (*g*) was then calculated from 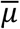 using Eq. [7].

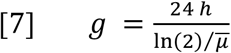

### Measurement of *proA\** copy number

The copy number of *proA\** was determined by qPCR of purified population genomic DNA. *gyrB* and *icd*, which remained at a single copy in the genome throughout the adaptation experiment, were used as internal reference genes. Details of the experimental procedure and data analysis are provided in the supplementary material.

### Whole-genome sequencing

Libraries were prepared from purified population genomic DNA using a modified Illumina Nextera protocol and multiplexed onto a single run on an Illumina NextSeq500 to produce 151-bp paired-end reads (Baym et al., 2015), giving a 60-130-fold coverage of the strain AM187 genome. Reads were trimmed using BBtools v35.82 (DOE Joint Genome Institute) and mapped using *breseq* v0.32.1 using the polymorphism (mixed population) option (Deatherage & Barrick, 2014).

### Growth rate measurements

Growth rates for each strain were calculated from growth curves measured in quadruplicate. Overnight cultures were grown in LB/kanamycin at 37 °C from glycerol stocks. Ten µL of each overnight culture was used to inoculate 4 mL of LB/kanamycin and the cultures were allowed to grow to mid-exponential phase (ODs_600_ 0.3-0.6) at 37 °C with shaking. The cultures were subjected to centrifugation at 4,000 × *g* for 10 min at room temperature and the pellets resuspended in an equal volume of PBS. The pellets were washed once more in PBS. The cells were diluted to an OD of 0.001 in Adaptation Medium and a 100 µL aliquot was loaded into each well of a 96-well plate. The plates were incubated in a Varioskan (Thermo Scientific) plate reader at 37 °C with shaking every 5 minutes for 1 minute. The absorbance at 600 nm was measured every 20 minutes for up to 500 hours. The baseline absorbance for each well (the average over several smoothed data points before growth) was subtracted from each point of the growth curve. Growth parameters (maximum specific growth, μ_max_; lag time, λ; maximum growth, A_max_) were estimated by non-linear regression using the modified Gompertz equation (Zwietering et al., 1990). Non-linear least-squares regression was performed in Excel using the Solver feature.

### Measurement of *argB* and *argH* gene expression by RT-qPCR

Overnight cultures were grown in LB/kanamycin at 37 °C from glycerol stocks. Ten µL of each overnight culture was used to inoculate 4 mL of LB/kanamycin and the cultures were grown to mid-exponential phase (OD_600_ 0.3-0.6) at 37 °C with shaking. Exponential phase cultures were centrifuged at 4,000 × *g* for 10 min and pellets resuspended in equal volume PBS. Pellets were washed once more in PBS. The cells were diluted to an OD of 0.001 in 4 mL of Adaptation Medium and grown to an OD_600_ of 0.2-0.3. Four 2-mL aliquots of culture were thoroughly mixed with 4 mL of RNAprotect Bacteria Reagent (Qiagen) and incubated at room temperature for 5 min before centrifugation at 4,000 × *g* for 12 min at room temperature. Pellets were frozen in liquid N_2_ and stored at −70 °C. Procedures for purification of RNA, reverse transcription, qPCR, and data analysis are described in the supplementary material.

### Measurement of ArgB and ArgH levels by mass spectrometry

Freezer stocks were used to streak each strain on LB/kanamycin. Four parallel 2-mL aliquots of LB/kanamycin were inoculated with individual colonies and the cultures were grown to mid-exponential phase at 37 °C with shaking. One mL of each culture was subjected to centrifugation at 16,000 × *g* for 1 min at room temperature. The cell pellets were resuspended in 1 mL PBS and washed twice more in PBS before resuspension and dilution to an OD of 0.001 in 5 mL of Adaptation Medium. Cultures were grown to an OD_600_ of 0.2-0.3 at 37 °C with shaking and then chilled on ice for 10 min before pelleting by centrifugation at 4,000 × *g* at 4 °C. Cell pellets were frozen in liquid N_2_ and stored at −70 °C.

Frozen cell pellets were thawed and lysed in 60 µL 50 mM Tris-HCl, pH 8.5, containing 4% (w/v) SDS, 10 mM TCEP and 40 mM chloroacetamide in a Bioruptor Pico sonication device (Diagenode) using 10 cycles of 30 seconds on, 30 seconds off, followed by boiling for 10 min, and then another 10 cycles in the Bioruptor. The lysates were subjected to centrifugation at 15,000 × *g* for 10 minutes at 20 °C and protein concentrations in the supernatant were determined by tryptophan fluorescence (Wiśniewski & Gaugaz, 2015). Ten µL of each sample (3-6 µg protein/µL) was digested using the SP3 method (C. S. Hughes et al., 2014). Carboxylate-functionalized speedbeads (GE Life Sciences) were added to the lysates. Addition of acetonitrile to 80% (v/v) caused the proteins to bind to the beads. The beads were washed twice with 70% (v/v) ethanol and once with 100% acetonitrile. Protein was digested and eluted from the beads with 15 µL of 50 mM Tris buffer, pH 8.5 with 1 µg endoproteinase Lys-C (Wako) for 2 hours with shaking at 600 rpm at 37 °C in a thermomixer (Eppendorf). One µg of trypsin (Pierce) was then added to the solution and incubated at 37 °C overnight with shaking at 600 rpm. Beads were collected by centrifugation and then placed on a magnet to more reliably remove the elution buffer containing the digested peptides. The peptides were then desalted using an Oasis HLB cartridge (Waters) according to the manufacturer’s instructions and dried in a speedvac.

Samples were suspended in 12 µL of 3% (v/v) acetonitrile/0.1% (v/v) trifluoroacetic acid and 500 ng of peptides were directly injected onto a C18 1.7 µm, 130 Å, 75 µm X 250 mm M-class column (Waters), using a Waters M-class UPLC. Peptides were eluted at 300 nL/minute using a gradient from 3% to 20% acetonitrile over 100 minutes into an Orbitrap Fusion mass spectrometer (Thermo Scientific). Precursor mass spectra (MS1) were acquired at a resolution of 120,000 from 380-1500 m/z with an AGC target of 2.0 × 10^5^ and a maximum injection time of 50 ms. Dynamic exclusion was set for 20 seconds with a mass tolerance of +/– 10 ppm. Precursor peptide ion isolation width for MS2 fragment scans was 1.6 Da using the quadrupole, and the most intense ions were sequenced using Top Speed with a 3-second cycle time. All MS2 sequencing was performed using higher energy collision dissociation (HCD) at 35% collision energy and scanning in the linear ion trap. An AGC target of 1.0 × 10^4^ and 35-second maximum injection time was used. Rawfiles were searched against the Uniprot *Escherichia coli* database using Maxquant version 1.6.1.0 with cysteine carbamidomethylation as a fixed modification. Methionine oxidation and protein N-terminal acetylation were searched as variable modifications. All peptides and proteins were thresholded at a 1% false discovery rate (FDR).

### Enzyme overexpression plasmids

*proA* and *proA\** were amplified from the genomes of *E. coli* BW25113 and AM187, respectively, and inserted into a linearized pET-46 vector backbone by Gibson assembly (Gibson et al., 2009). A sequence encoding a 6×His-tag followed by a 2×Val-linker was incorporated at the N-terminus of each protein. The *proA*\*\* expression plasmid was constructed using the Q5 Site-Directed Mutagenesis Kit (NEB).

*argC* was cloned into a pTrcHisB vector backbone. The final plasmid included an N-terminal 6×His-tag followed by a Gly-Met-Ala-Ser linker and with Met1 of ArgC removed.

The *argD* and *argI* expression plasmids from the ASKA collection (Kitagawa et al., 2005) were used for expression of *N*-acetylornithine aminotransferase and ornithine transcarbamoylase, respectively. In each case, the expression plasmid included a sequence encoding an N-terminal 6×His-tag followed by a Thr-Asp-Pro-Ala-Leu-Arg-Ala linker.

Wild-type *carAB* was amplified from the genome of AM187 and inserted into a linearized pCA24N vector backbone by Gibson assembly (Gibson et al., 2009). The final construct included an N-terminal 6×His-tag on CarA followed by a Thr-Asp-Pro-Ala-Leu-Arg-Ala linker. CPS mutant plasmids were constructed using the Q5 Site-Directed Mutagenesis Kit (NEB).

The correct sequences for all constructs were confirmed by Sanger sequencing.

### Protein purification

Wild-type and variant ProAs were expressed in strain AM209 (BL21(DE3) *argC*::*kan*^*r*^ *proA*::*cat*) to enable purification in the absence of wild-type ProA and ArgC. Carbamoyl phosphate synthetase (CPS) consists of a stable complex between CarA and CarB. Thus, in order to purify CPS, both *carA* and *carB* were co-expressed on the same plasmid with a His-tag on CarA. Wild-type and variant CPSs were expressed in strain AM267 (BL21 *carAB*::*kan*^*r*^) to enable purification in the absence of wild-type CPS.

Enzymes were expressed and purified using the following protocol with minor variations between the proteins. A small scraping from the glycerol stock of each expression strain was used to inoculate LB plus appropriate antibiotics. The cultures were grown overnight with shaking at 37 °C. Overnight cultures were diluted 1:100 into 500 mL-2 L of LB plus appropriate antibiotic and grown with shaking at 37 °C. Once the OD_600_ reached 0.5-0.9, IPTG was added to a final concentration of 0.5 mM. Growth was continued at 30 °C for 5 hrs while shaking. Cells were harvested by centrifugation at 5,000 × *g* for 20 min at 4 °C. Cell pellets were stored at −70 °C until protein purification.

Frozen cell pellets were resuspended in 5× the cell pellet weight of ice-cold 20 mM sodium phosphate, pH 7.4, containing 300 mM NaCl and 10 mM imidazole. Fifty µL of protease inhibitor cocktail (Sigma-Aldrich, P8849) was added for each gram of cell pellet. Lysozyme was added to a final concentration of 0.2 mg/mL and the cells were lysed by probe sonication (20 sec of sonication followed by 30 sec on ice, repeated 3 times). Cell debris was removed by centrifugation at 18,000 × *g* for 20 min at 4 °C. The soluble fraction was then loaded onto 1 mL or 3 mL HisPur Ni-NTA Spin Columns (Thermo Scientific) and His-tagged protein was purified according to the manufacturer’s protocol. If the soluble fraction was larger than the column volume, the initial binding step was repeated before washing to accommodate all the soluble fraction. Bound protein was eluted with one column volume of 20 mM sodium phosphate, pH 7.4, containing 300 mM NaCl and increasing amounts of imidazole (100 mM, 250 mM, and finally 500 mM). Two separate elutions were performed with 500 mM imidazole. Fractions containing the protein of interest were pooled and dialyzed overnight against 6-12 L of exchange buffer at 4 °C. (ProA and ArgC were dialyzed against 20 mM potassium phosphate, 20 mM DTT, pH 7.5. *N*-acetylornithine aminotransferase was dialyzed against 20 mM potassium phosphate, pH 7.5. CPS was dialyzed against 100 mM potassium phosphate, pH 7.6. Ornithine transcarbamoylase was dialyzed against 20 mM Tris-acetate, pH 7.5.) Protein purity was assessed by SDS-PAGE and concentration measured using the Qubit protein assay kit with a Qubit 3.0 fluorometer (Invitrogen). Purified protein was stored at 4 °C for short-term storage, and frozen in liquid nitrogen and stored at −70 °C for long-term storage.

### GSA and NAGSA dehydrogenase assays

The native and neo-ArgC activities of ProA were assayed in the reverse direction (dehydrogenase reaction) because the lability of the forward substrates γ-glutamyl phosphate and *N*-acetylglutamyl phosphate make them difficult to purify. The change in the dehydrogenase activity due to a mutation is proportional to the change in the reductase activity according to the Haldane relationship (Haldane, 1930; McLoughlin & Copley, 2008).

Assaying ProA’s dehydrogenase activity using γ-glutamyl semialdehyde (GSA) and *N*-acetylglutamyl semialdehyde (NAGSA) as substrates is complicated by the equilibrium of GSA and NAGSA with their hydrated forms, as well as GSA’s intramolecular cyclization to form pyrroline-5-carboxylate (P5C) (Bearne & Wolfenden, 1995; Mezl & Knox, 1976). In order to measure the concentration of the free aldehyde form of these substrates, we mixed 15 µM ProA or ArgC with 2 mM “GSA” (including the hydrate and P5C) or 2 mM “NAGSA” (including the hydrate), respectively, in a solution containing 100 mM potassium phosphate, pH 7.6, and 1 mM NADP^+^ and measured the burst in NADPH production (Khanal et al., 2015). The concentrations of GSA+P5C+hydrate or NAGSA+hydrate were determined using the *o*-aminobenzaldehyde assay (Albrecht et al., 1962; Mezl & Knox, 1976). The absorbance at 340 nm due to formation of NADPH exhibited a burst followed by a linear phase when measured for 60 seconds. We assume that the burst corresponds to reduction of the free aldehyde form of GSA or NAGSA and the rate of the linear phase is determined by the conversion of the hydrate (and P5C in the case of GSA) to the free aldehyde. We calculated the magnitude of the burst by fitting either all of the data or the linear portion of the data to one of the following equations.

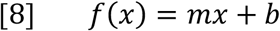

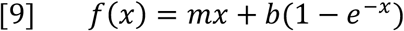

where *x* is time in seconds, *m* is the slope of the linear phase, and *b* is the magnitude of the burst and thus proportional to the starting concentration of the free aldehyde form of the substrate. In the case of the linear fit, only the linear portion of the A_340_ data was used. Eq. [8] was used to calculated NAGSA free aldehyde concentration because the exponential equation did not fit the data well. Eq. [9] was used to calculated GSA free aldehyde concentration. We repeated the assay three times and averaged the magnitude of the burst to calculate free aldehyde concentrations for solutions of GSA and NAGSA (under these buffer and temperature conditions) of 4.5% and 4.2% of the total concentration of free aldehyde + hydrate (+ P5C for GSA), respectively.

GSA and NAGSA dehydrogenase activities were measured by monitoring the appearance of NADPH at 340 nm in reaction mixtures containing 100 mM potassium phosphate, pH 7.6, 1 mM NADP^+^, varying concentrations of NAGSA or GSA, and catalytic amounts of ProA, ProA*, and ProA**. All kinetic measurements were done at 25 °C. Values for *K*_*M*_ refer to the concentration of the free aldehyde form of the substrate.

### Assays for carbamoyl phosphate synthetase activity and allosteric regulation

Kinetic assays for carbamoyl phosphate synthetase (CPS) were carried out with minor modifications of the methods described in Pierrat and Raushel 2002. The rate of ATP hydrolysis was measured at 37 °C by coupling production of ADP to oxidation of NADH using pyruvate kinase, which converts ADP and PEP to ATP and pyruvate, and lactate dehydrogenase, which reduces pyruvate to lactate. Loss of NADH was monitored at 340 nm. Reaction mixtures consisted of 50 mM HEPES, pH 7.5, containing 10 mM MgCl_2_, 100 mM KCl, 20 mM potassium bicarbonate, 10 mM L-glutamine, 1 mM PEP, 0.2 mM NADH, saturating amounts of pyruvate kinase and lactate dehydrogenase (Sigma-Aldrich, P0294), and varying amounts of ATP (0.01 to 8 mM). Reactions were initiated by adding 0.2 µM CPS to a final concentration of 0.2 µM. The effects of UMP and ornithine were measured under the same reaction conditions but with a fixed ATP concentration of 0.2 mM and varying concentrations of either UMP or ornithine.

Carbamoyl phosphate production was measured with minor modifications of previously described procedures (Snodgrass & Parry, 1969; Stapleton et al., 1996). Formation of carbamoyl phosphate by CPS was coupled with formation of citrulline by ornithine transcarbamoylase; citrulline forms a yellow complex (ε_464_ = 37800 M^-1^ cm^-1^ (Snodgrass & Parry, 1969)) when mixed with diacetyl monoxime and antipyrine. Reaction mixtures consisted of 50 mM HEPES, pH 7.5, 10 mM MgCl_2_, 100 mM KCl, 20 mM potassium bicarbonate, 10 mM L-glutamine, 4 mM ATP, 10 mM L-ornithine, and 0.7 µM ornithine transcarbamoylase. Reactions (0.25 mL) were initiated by adding CPS at a final concentration of 0.2 µM. After incubation for 2.5 min at 37 °C, reactions were quenched by addition of 1 mL of a solution consisting of 25% concentrated H_2_SO_4_, 25% H_3_PO_4_ (85%), 0.25% (w/v) ferric ammonium sulfate, and 0.37% (w/v) antipyrine, followed by addition of 0.5 mL of 0.4% (w/v) diacetyl monoxime/7.5% (w/v) NaCl. The quenched reaction mixtures were placed in a boiling water bath for 15 min before measurement of OD_464_. Control reactions contained all components except CPS.

### RNA structure prediction

RNA secondary structures for *argB* mRNAs were predicted using CLC Main Workbench 8.1. The entire intergenic region between *kan*^*r*^ and *argB* plus the first 33 nucleotides of *argB* were included in the structure prediction. The first 33 nucleotides were included because an mRNA-bound ribosome prevents another ribosome from binding to the mRNA until it has moved past the first 33 nucleotides (Steitz, 1969). Thus, at least the first 33 nucleotides are available for folding with the upstream region when a mRNA is being translated.

### Calculation of RNA folding times

Folding times for a 63-nucleotide region (30 nt downstream and 30 nt upstream of the *argB* start codon) surrounding the start codon of *argB* mRNAs were calculated using the Kinfold program (v1.3) from the ViennaRNA v2.4.11 package (Wolfinger et al., 2004). Kinfold utilizes a Monte Carlo algorithm to calculate the folding time of each RNA sequence to the lowest free energy structure. We simulated 500 folding trajectories for each structure.

## Acknowledgements

This work was supported by the NASA Astrobiology Institute (NNA15BB04A). We thank Craig Joy (University of Colorado Boulder, Physics Department) and Chris Takahashi (University of Washington) for help building the turbidostat.

## Funding

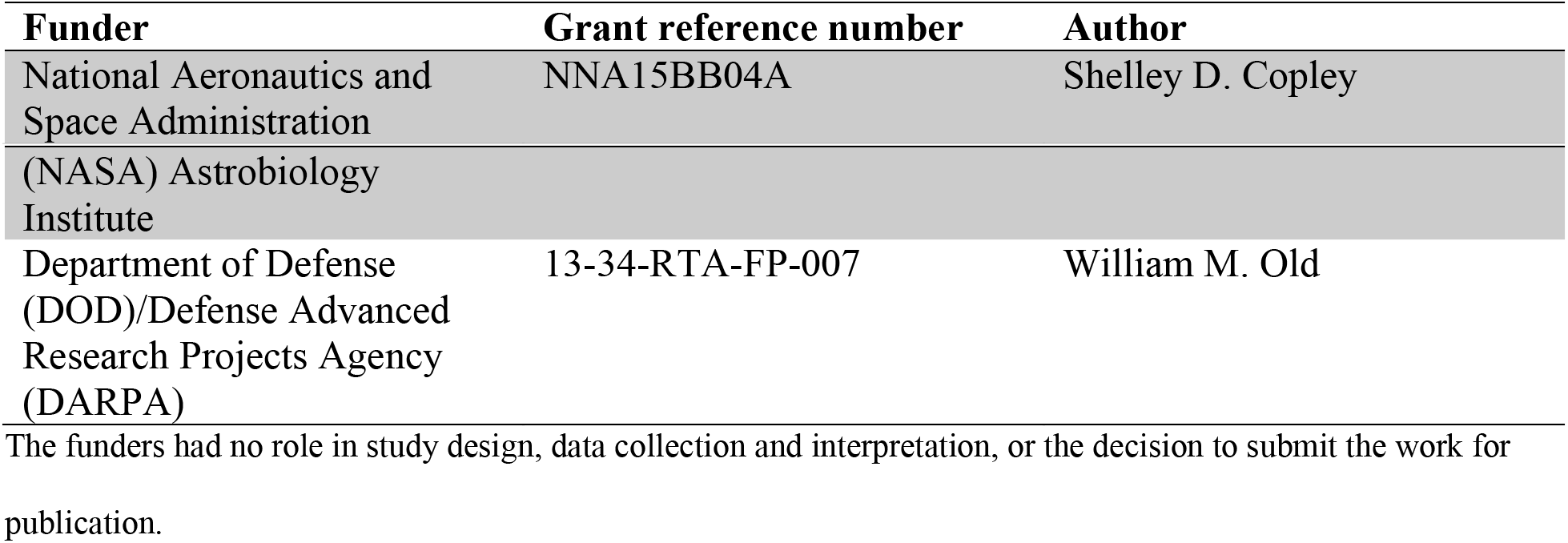

## Additional files

**Supplementary file 1.** Mutations found during the adaptation experiment. Sheet 1: mutations that were present at frequencies greater than 30% or that appeared in different populations. Sheet 2: genome positions of the amplified regions surrounding *proA\**. Sheet 3: other amplified or deleted genomic regions in the adapted populations.

**Supplementary file 2.** Supplementary Materials and Methods.

## Data availability

The genome sequence of *E. coli* strain AM187 used in this study has been deposited to NCBI GenBank under accession number CP037857.

## Supplementary file 2: Supplementary Materials and Methods

**Table S1.** Plasmids used in this study.

| Plasmid | Use/description | Source/Ref |
| --- | --- | --- |
| pSLTS | scarless genome editing/arabinose-inducible $\lambda$ Red recombinase, anhydrotetracycline-inducible meganuclease I-SceI, temperature-sensitive origin of replication, ampicillin resistance | (Kim et al., 2014) |
| pSIM5 | introduction of linear DNA fragments into the genome/heat-inducible $\lambda$ Red recombinase, temperature sensitive origin of replication, chloramphenicol resistance | (Datta et al., 2006) |
| pSIM27 | introduction of linear DNA fragments into the genome/heat-inducible $\lambda$ Red recombinase, temperature sensitive origin of replication, tetracycline resistance | (Datta et al., 2006) |
| pAM053 | Cas9-mediated genome editing/constitutively-expressed *cas9*, heat-inducible $\lambda$ Red recombinase, temperature-sensitive origin of replication, chloramphenicol resistance | This study |
| pET-46 | backbone for IPTG-inducible expression of wild-type and mutant ProAs/T7 promoter upstream of 6xHis-tag, ampicillin resistance | Novagen |
| pCA24N | backbone for IPTG-inducible expression of wild-type and mutant carbamoyl phosphate synthetases/chloramphenicol resistance | (Kitagawa et al., 2005) |
| pCA24N-*argI* | IPTG-inducible expression of ornithine transcarbamoylase/chloramphenicol resistance | (Kitagawa et al., 2005) |
| pCA24N-*argD* | IPTG-inducible expression of *N*-acetylornithine aminotransferase/chloramphenicol resistance | (Kitagawa et al., 2005) |
| pAM068 | expresses the Cas9 guide RNA that targets the upstream region of *argB* for introduction of the 58 bp deletion upstream of *argB*/temperature-sensitive origin of replication, ampicillin resistance | This study |
| pAM100 | expresses the Cas9 guide RNA that targets *rph* for introduction of the 82 bp deletion upstream of *pyrE*/temperature sensitive origin of replication, ampicillin resistance | This study |

**Table S2.**
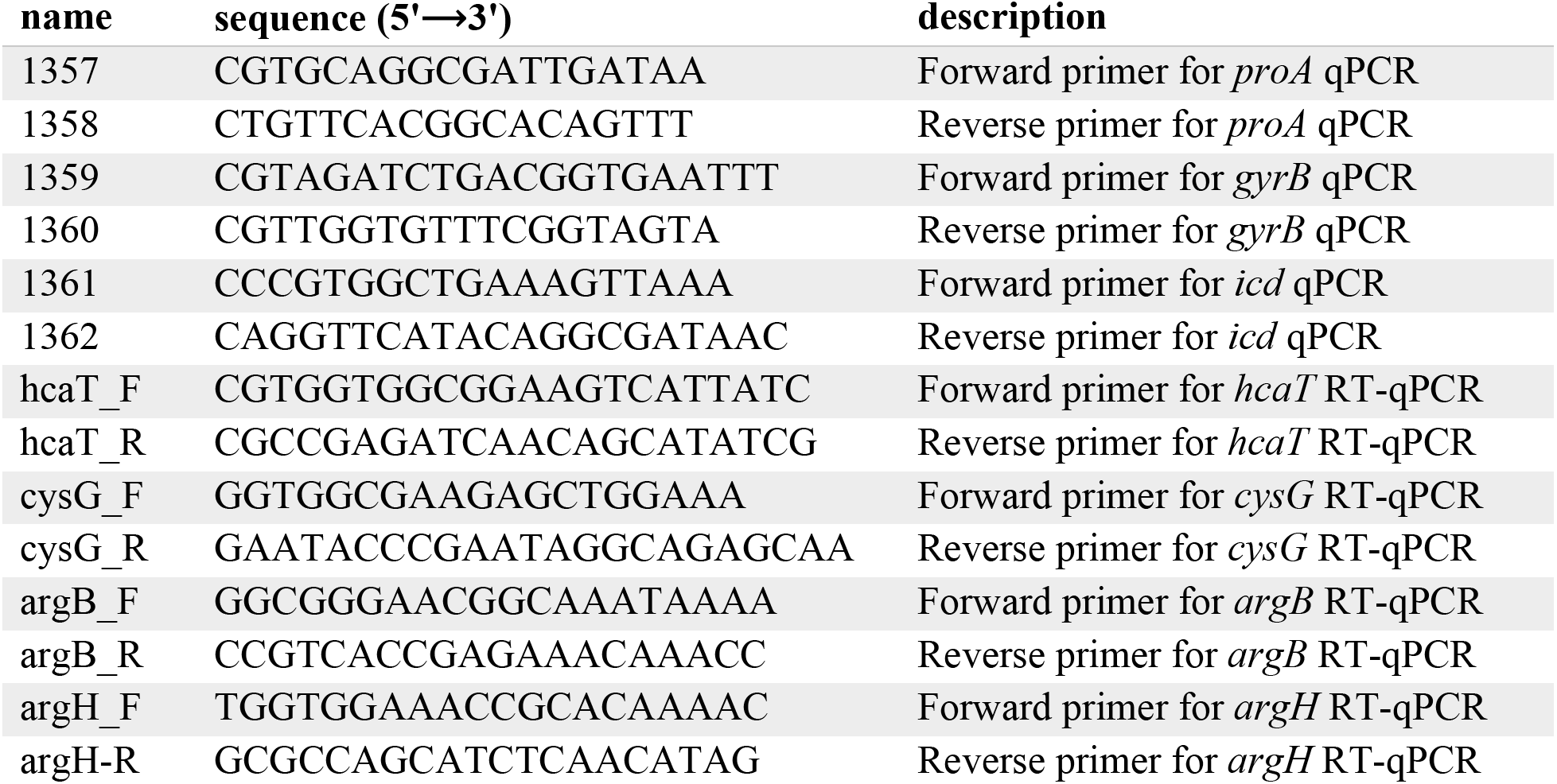
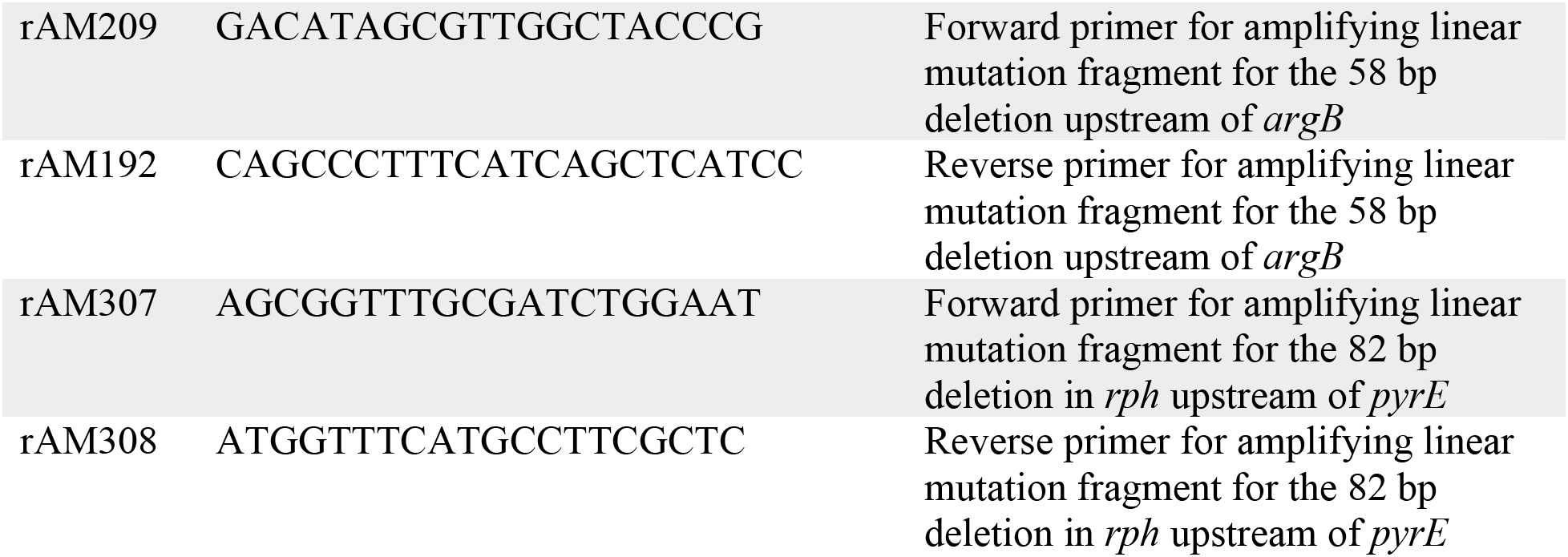
Primers used in this study.

**Table S3.**
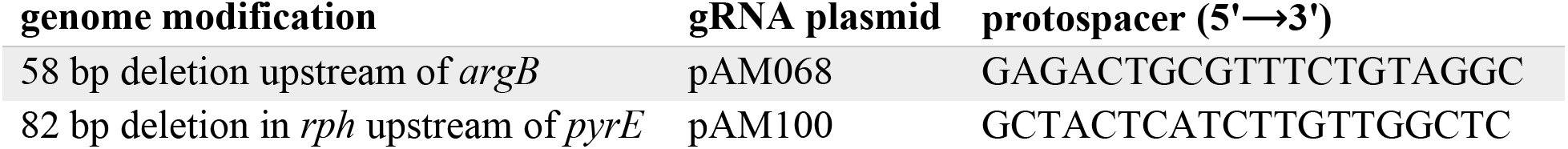
Protospacers used for Cas9-mediated scarless genome editing.

**Table S4.**
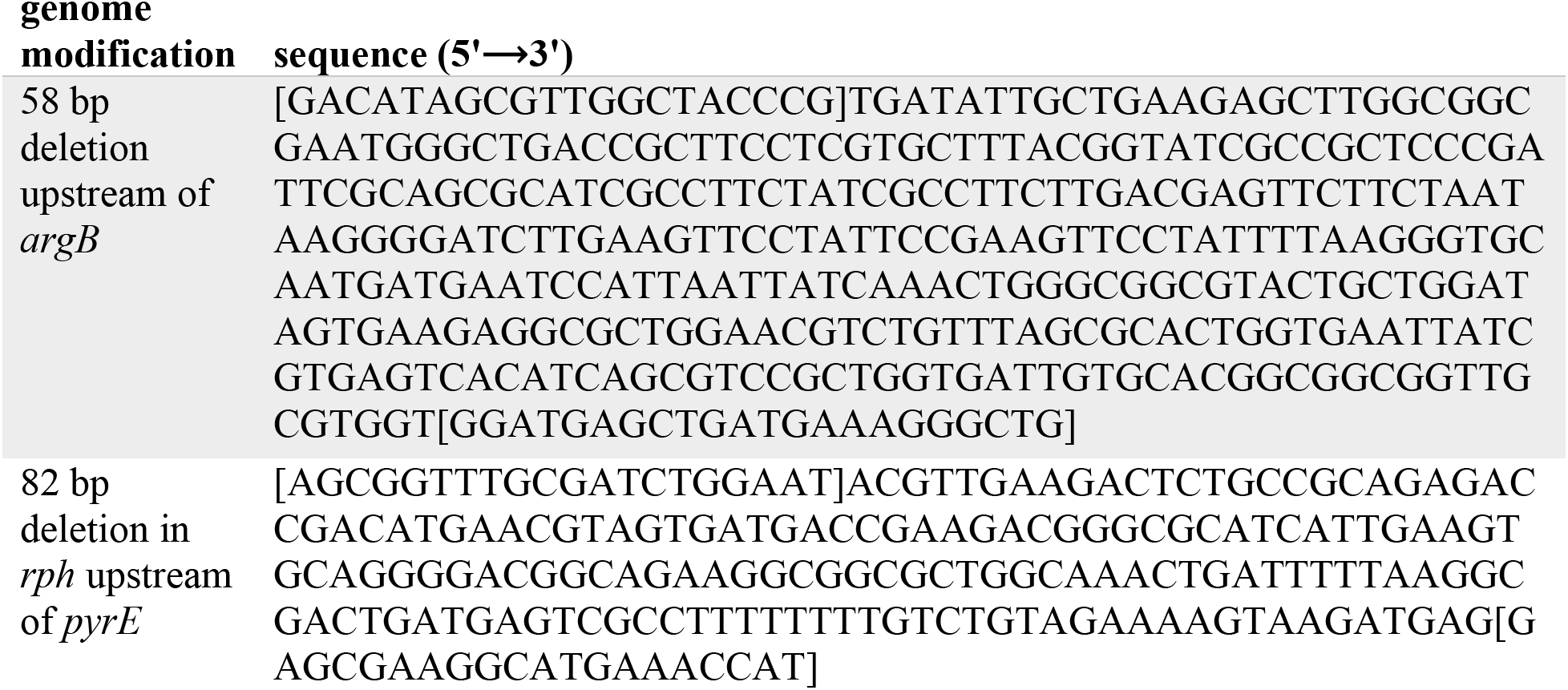
Mutation cassettes used for Cas9-mediated scarless genome editing. Brackets represent the primer annealing regions for amplifying the mutation cassettes.

### Genome editing

Strain AM187 was first transformed with a helper plasmid (pAM053, Table S1) encoding *cas9* under the control a weak constitutive promoter (pro1 from Davis, Rubin, and Sauer 2011), *λ* Red recombinase genes (*exo*, *gam*, and *bet*) under the control of a heat-inducible promoter, and a temperature-sensitive origin of replication (Datta et al., 2006). These cells were grown to an OD_600_ of 0.2-0.4 at 30 °C and then incubated at 42 °C with shaking for 15 min to induce expression of the *λ* Red recombinase genes. The cells were immediately subjected to electroporation with 100 ng of either pAM068 or pAM100, which encode guide RNAs targeting 20-nucleotide sequences upstream of *argB* or in *rph*, respectively (Table S3), and 450 ng of a linear homology repair template that will introduce the desired deletion into the genome (Table S4). (Linear homology repair templates were amplified from genomic DNA of clones isolated during the adaptation experiment that contained the desired deletions and and the PCR fragments were gel-purified. Primers used to generate the linear DNA mutation fragments are listed in Table.) The cells were allowed to recover at 30 °C for 2-3 hours before being spread onto LB/ampicillin plates. Surviving colonies contained the guide RNA plasmid and the desired deletion, as confirmed by Sanger sequencing. pAM053 and the guide RNA plasmids, both of which have temperature-sensitive origins of replication, were cured by growth of individual colonies at 37 °C.

Strain AM209 was constructed from *E. coli* BL21(DE3) for expression of wild-type and mutant ProAs. We deleted *argC* and *proA* to ensure that any activity measured during *in vitro* assays was not due to trace amounts of ArgC or wild-type ProA. To accomplish these deletions, we amplified and gel-purified DNA fragments containing antibiotic resistance genes (kanamycin and chloramphenicol for deletion of *argC* and *proA*, respectively) flanked by 200-400 bp of sequences homologous to the upstream and downstream regions of either *argC* or *proA*. *E. coli* BL21(DE3) cells containing pSIM27 (Datta et al., 2006) – a vector containing heat-inducible *λ* Red recombinase genes – were grown in LB/tetracycline to an OD of 0.2-0.4 and then incubated in a 42 °C shaking water bath for 15 min to induce expression of *λ* Red recombinase genes. The cells were then immediately subjected to electroporation with 100 ng of the appropriate linear DNA mutation cassette. Successful transformants were selected on either LB/kanamycin or LB/chloramphenicol plates.

Strain AM267 was constructed from *E. coli* BL21(DE3) for expression of wild-type and mutant carbamoyl phosphate synthetases (CPS). We deleted *carAB* to ensure that any activity measured during *in vitro* assays was not due to trace amounts of wild-type CPS. To accomplish the deletion, we amplified and gel-purified a DNA fragment containing the kanamycin resistance gene flanked by 40 bp of sequence homologous to the upstream and downstream regions of either *argC* or *proA*. *E. coli* BL21 cells containing pSIM5 (Datta et al., 2006) – a vector carrying heat-inducible *λ* Red recombinase genes – were grown in LB/chloramphenicol to an OD of 0.2-0.4 and then incubated in a 42 °C shaking water bath for 15 min to induce expression of *λ* Red recombinase genes. The cells were then immediately subjected to electroporation with 100 ng of the appropriate linear DNA mutation cassette. Successful transformants were selected on either LB/kanamycin plates.

### Measurement of *proA\** copy number

The primer sets used for each gene are listed in Table S2. PowerSYBR Green PCR master mix (Thermo Scientific) was used according to the manufacturer’s protocol. A standard curve using variable amounts of AM187 genomic DNA was run on every plate to calculate efficiencies for each primer set. Primer efficiencies were calculated with the following equation:

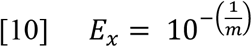

where *E* is the efficiency of primer set *x*, and *m* is the slope of the plot of C_t_ vs. starting quantity for the standard curve. *proA\** copy number was then calculated with the following equation (Hellemans et al., 2007):

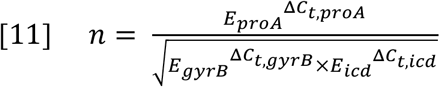

where *n* is the *proA\** copy number, and *∆C*_*t,x*_ is the difference in Ct’s measured during amplification of AM187 and sample genomic DNA with primer set *x*.

### Measurement of *argB* and *argH* gene expression by RT-qPCR

RNA was purified using the Invitrogen PureLink RNA Mini Kit according to the manufacturer’s protocol. The cell lysate produced during the PureLink protocol was homogenized using the QIAShredder column (Qiagen) prior to RNA purification. After RNA purification, each sample was treated with TURBO DNase (Invitrogen) according to the manufacturer’s protocol. Reverse transcription (RT) was performed with 250-600 ng of RNA using SuperScript IV VILO (Invitrogen) master mix according to the manufacturer’s protocol.

qPCR of cDNA was performed to measure the fold-change in expression of *argB* and *argH* in mutant strains compared to that in AM187. *hcaT* and *cysG* were used as reference genes (Zhou et al., 2011). The primer sets used for each gene are listed in Table S2. A standard curve using variable amounts of *E. coli* BW25113 genomic DNA was run to calculate the primer efficiencies for each primer set. Fold-changes in expression of *argB* and *argH* were calculated as described above for calculations of *proA\** copy number.

